# Conformational ensembles of flexible multidomain proteins: How close are we to accurate and reliable predictions?

**DOI:** 10.64898/2026.02.24.707687

**Authors:** Santiago Rodriguez, Aurélie Fournet, Simon Bartels, Mátyás Pajkos, Ilinka Clerc, Laure Carrière, Aurélien Thureau, Cédric Montanier, Claire Dumon, Frédéric Allemand, Juan Cortés, Pau Bernadó

## Abstract

Multidomain proteins connected by flexible linkers populate conformational ensembles that are challenging to characterize using conventional structural biology methods. In domain–linker–domain (DLD) proteins, linker-mediated inter-domain relative positions and orientations are functionally relevant, yet their dynamical behavior in solution normally remain poorly described. Small-angle X-ray scattering (SAXS) provides ensemble-averaged structural information for such systems; however, coupling with computational modeling is required to accurately describe the dynamic behavior of this family of proteins in solution. Here, we present a systematic evaluation of five ensemble-generation strategies applied to a set of eighteen proteins sharing the same two globular domains, connected by naturally occurring linkers of varying length and composition. Modeling methods based on different underlying principles are compared by assessing their agreement to experimental SAXS data, showing a large disparity and systematic structural biases among them. Furthermore, for each approach, we examine the effect of refinement against SAXS restraints and assess its capacity to describe the experimental data, as well as the induced biases in global dimensions and inter-domain distance distributions. This analysis underlines the importance of the initial conformational pool for deriving experimentally compatible ensembles. Overall, this work provides a high-quality benchmark for SAXS-driven ensemble modeling of flexible, multidomain proteins and establishes a framework for the critical interpretation of solution scattering data in systems with pronounced conformational heterogeneity.

## 1 Introduction

Many proteins exhibit multimodular architectures in which structured domains are connected by flexible or intrinsically disordered linkers [1, 2]. These domain–linker–domain (DLD) proteins are widespread across all kingdoms of life and play key roles in numerous biological processes [3–6]. The presence of flexible linkers enables functional adaptability by allowing domains to sample a wide range of relative orientations and distances, facilitating molecular recognition, catalysis, and regulation [1, 6]. At the same time, this conformational heterogeneity poses significant challenges for structural characterization and quantitative modeling.

A prominent example of DLD architectures is found in enzymes involved in biomass degradation, where catalytic glycosyl hydrolase (GH) domains are often connected to carbohydrate-binding modules (CBMs) through flexible linkers (Figure 1). In these systems, the linker length and composition modulate the spatial relationship between catalytic and binding domains, directly influencing substrate targeting, enzymatic efficiency and overall function [7, 8]. As a result, GH–CBM chimeras have become important targets in enzyme engineering and biotechnology, motivating a deeper understanding of how linker properties shape their conformational ensembles in solution [7–9].

**Figure 1:**
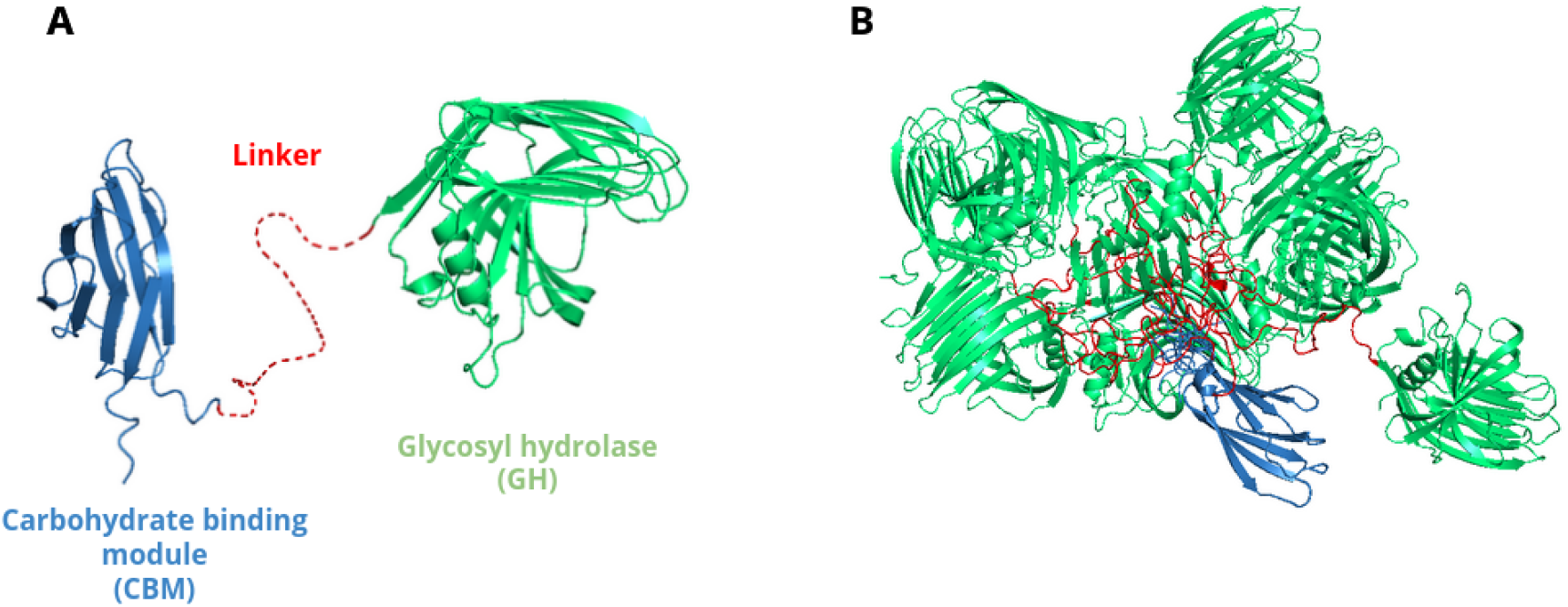
(A) Generic structure of the DLD proteins studied in this work. Cartoon representations correspond to the glycosyl hydrolase (GH; in green) and carbohydrate-binding module (CBM; in blue) domains, while the red dashed line represents the connecting linker, which varies in both length and composition across the different DLD systems. (B) Generic ensemble representation of a DLD protein. The relative pose (position and orientation) between the CBM and GH domains is defined by the linker.

Traditional high-resolution structural biology techniques such as X-ray crystallography and cryo-electron microscopy have inherent limitations when applied to highly flexible or partially disordered proteins [10,11]. Although more adapted to the conformational plasticity of DLD [12–14], Nuclear Magnetic Resonance (NMR) often faces severe spectral overlap with large domains. As a consequence, flexible linkers often remain unresolved and the relative positions of the domains are unknown. In this context, small-angle X-ray scattering (SAXS) has emerged as a powerful technique that probes proteins in solution and reports on their ensemble-averaged structural properties [15, 16]. However, SAXS data are intrinsically low-resolution and do not provide unique structural solutions, making their interpretation strongly dependent on computational modeling [17–19]. Consequently, the generation of realistic conformational ensembles from sequence or structural information has become a central component in the analysis of SAXS data for flexible proteins [20, 21].

Molecular dynamics (MD) simulations using all-atom physics-based force field models have been widely used for the investigation of globular proteins. Although molecular force-fields and water models have dramatically improved in the recent years [22, 23], and new protocols have been developed to enhance the conformational sampling [24], MD simulations remain still limited to study large, highly flexible proteins [25]. Alternatively, a wide range of computational approaches has been developed for this purpose [26], including physics-based coarse-grained models [27, 28], structure-encoding libraries [29–31] and, more recently, methods derived from AlphaFold-based frameworks [32]. These approaches have demonstrated encouraging performance in various contexts, including community-wide benchmarks such as CASP [33]. Nevertheless, their general applicability and reliability for highly flexible, multidomain proteins remain uneven, and systematic evaluations focused on DLD architectures are still scarce.

Several key questions therefore remain open: How accurately do current ensemble generation methods reproduce experimental SAXS profiles for DLD proteins? To what extent and in which conditions can refinement against SAXS data improve the agreement between predicted and experimental observables? And do ensembles generated by different methods converge toward similar conformational descriptions after refinement, or do method-dependent differences persist? Addressing these questions is essential for establishing confidence in ensemble modeling strategies and for defining best practices in the interpretation of SAXS data.

In this work, we address these issues by systematically evaluating multiple computational approaches for generating conformational ensembles of GH11–CBM chimeras connected by linkers of diverse length and composition and for which we have recorded high-quality SAXS data. These profiles constitute an interesting benchmark set to assess the capacity of current ensemble generation methods to reproduce experimental SAXS profiles for DLD proteins. For each method, the quality of the resulting ensemble is evaluated by directly comparing with the experimental profile. Moreover, the structural impact of ensemble refinement against SAXS data is quantitatively assessed. By refining ensembles derived from distinct starting pools and comparing their resulting structural properties, we examine the consistency and robustness of the inferred conformational descriptions.

Beyond the specific protein family studied here, our results provide insights into the strengths and limitations of current ensemble modeling strategies for DLD proteins. Such understanding is critical not only for advancing fundamental studies of protein conformational behavior, but also for the rational design and optimization of multimodular enzymes with applications in biotechnology and industrial biocatalysis.

## 2 Results

### 2.1 Protein system and linker diversity

For our benchmark set, we constructed a series of chimeric proteins consisting of the catalytic xylanase GH11 domain from *Neocallimastix patriciarum* (UniProt: P29127; PDB: 2C1F) connected to a CBM from *Cellulomonas fimi* (UniProt: P54865; PDB: 2XBD) through natural linker sequences extracted from the CAZy database [34]. From more than one thousand available protein fragments encompassing GH11–linker-CBM domains, we selected 18 linker sequences that were inserted between both globular domains for subsequent structural characterization. The selection was guided by sequence-based descriptors capturing key features of linker diversity, specially length and amino acid composition. This strategy ensured that sequences with diverse charge distribution, proline and glycine content or low complexity segments were represented in both short (10 residues) and long (>30 residues) linkers.

Table 1 presents the sequences and a set of descriptors of the selected linkers for the construction of the chimeras, which were named from DLD1 to DLD18. Physicochemical properties were computed using localCIDER [35]. The linkers sampled a broad range of sizes, from 10 to 88 residues. In terms of composition, while low complexity linkers were rich in glycines (DLD1, 4 and 8), prolines (DLD10 and 15), serines (DLD9 and 13) or asparagines (DLD16), others presented less compositional bias (DLD3, 7, 17 and 18). In terms of electrostatic charges, the vast majority of linkers were neutral, and only DLD3 (-2), DLD11 (+5) and DLD17 (-9) displayed a moderate global net charge. Thus, this panel of proteins captured the natural variability of GH11-associated linkers while providing a well-defined system with identical folded domains.

**Table 1:**
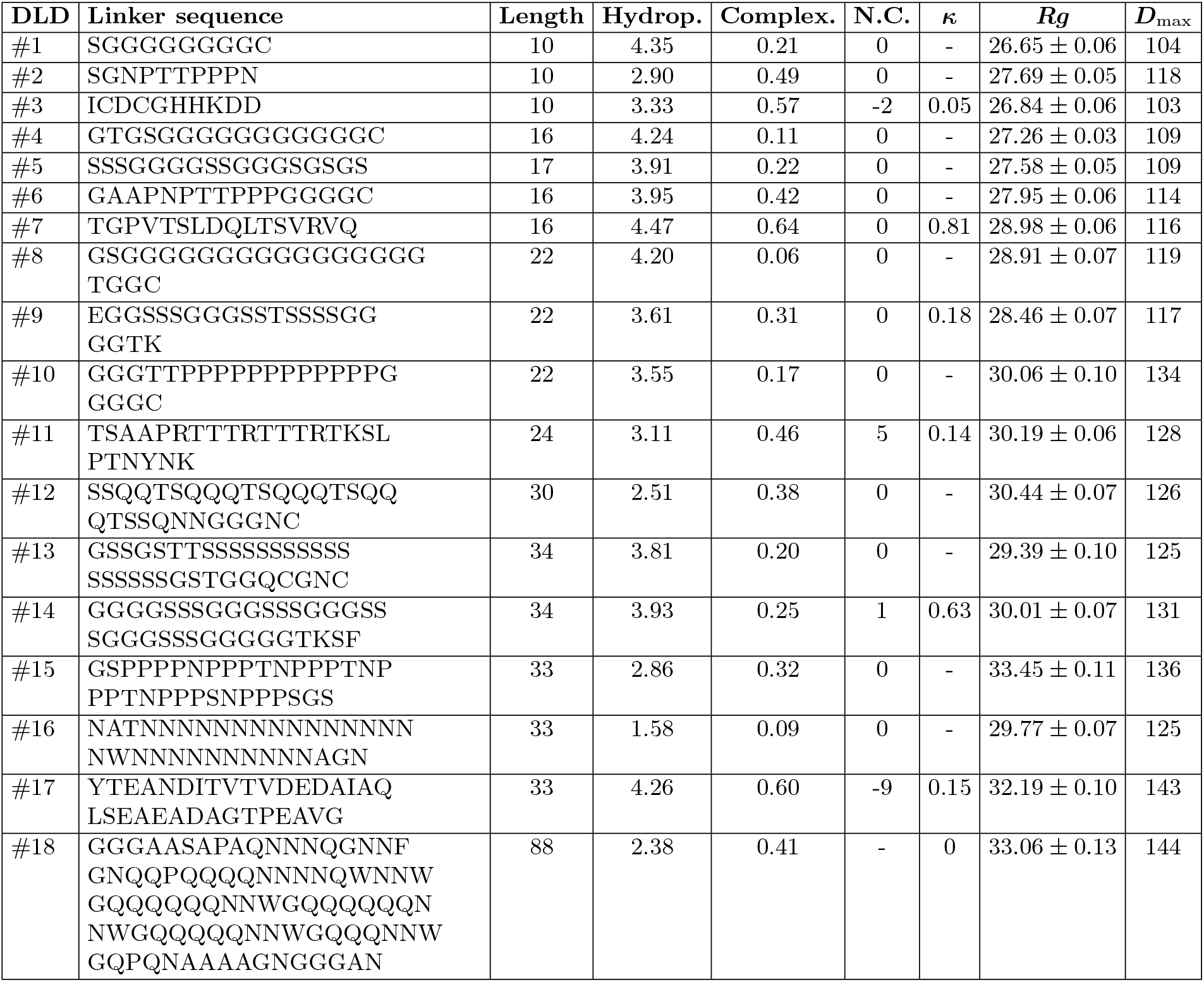
Features of the selected linkers from the CAZy database. Hydropathy (Hydrop.) indicates the relative hydrophobicity of a sequence; higher values correspond to higher hydrophobicity. Complexity (Complex.) estimates the local compositional complexity of a sequence; lower values indicate simpler, low-complexity regions. N.C. indicates the total charge of the sequence. *κ* measures charge patterning in proteins; higher values indicate more segregated charges and a tendency toward compact conformations. *Rg* and *D*_max_ values, in Å, were derived from the analyses of the SAXS experiments.

### 2.2 Small-angle X-ray Scattering shows the flexibility of the DLD benchmark set

A plasmid was designed in order to facilitate the subsequent cloning of the 18 linkers between the GH and CBM domains (see Materials and Methods for details). The resulting chimeras were overexpressed in *Escherichia coli* and purified following previously developed procedures for this family of proteins (see methods section for details) [36, 37]. The purified proteins were submitted to size-exclusion chromatography coupled to small-angle X-ray scattering (SEC-SAXS) measurements. The concentrations used for the injections are shown in Figure S1.

In all cases, the SEC profile showed a single peak that was treated using standard procedures to derive a final SAXS profile for subsequent analyses. All the curves displayed an excellent signal to noise and a complete absence of concentration-dependent phenomena, such as oligomerization and interparticle interactions (Fig. 2 and S1 in S.I.). These features underline the quality of this benchmark set to test present protein disorder modelling tools (see below).

**Figure 2:**
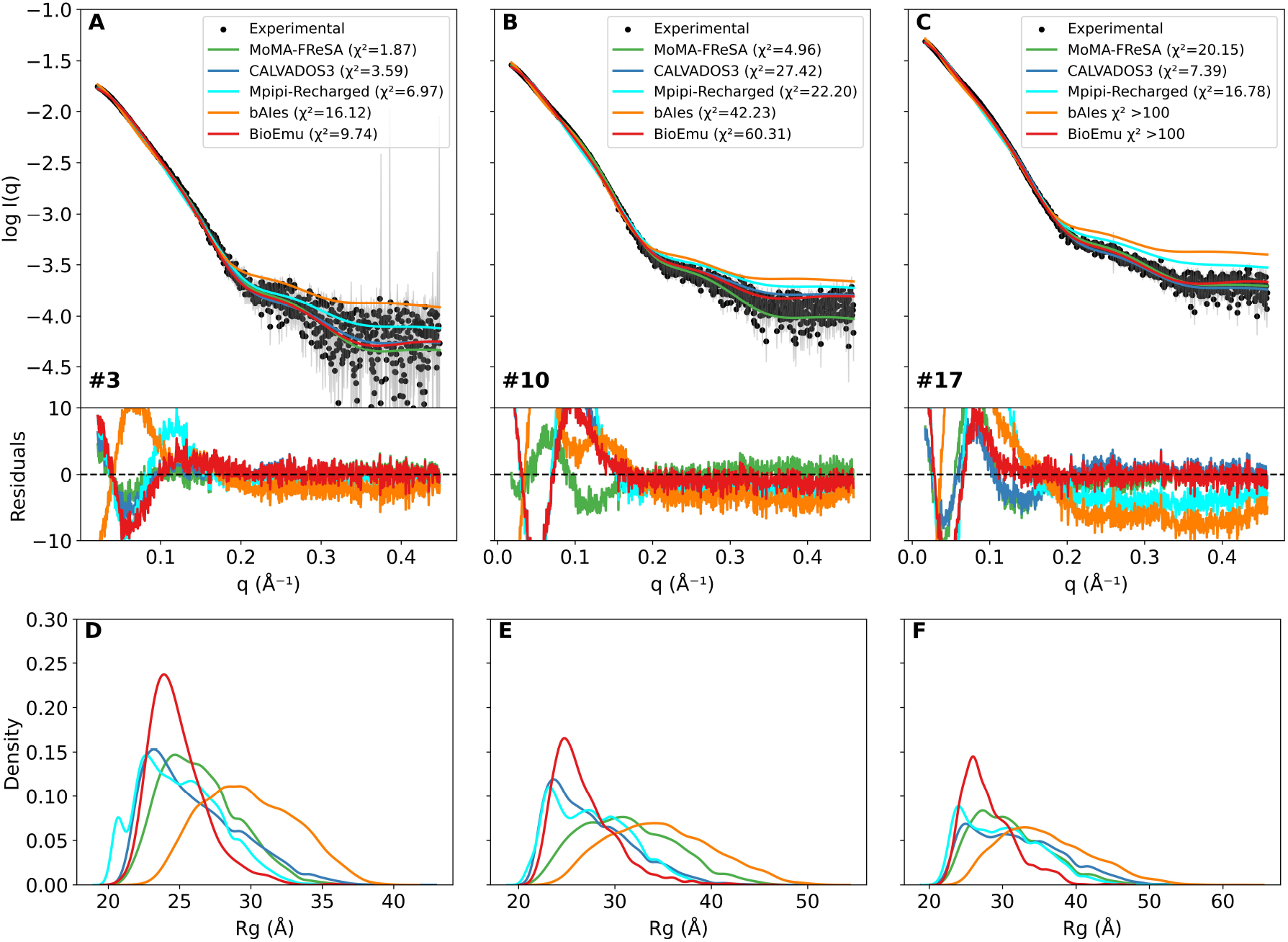
SAXS intensity profiles (black) for DLD3 (A), DLD10 (B) and DLD17 (C) compared with the averaged scattering profiles computed from the complete ensembles modelled with MoMA-FReSa (green), CALVADOS3 (blue), Mpipi-Recharged (cyan), bAIes (orange) and BioEmu (red). *χ*^2^-values are shown in each panel. Point-by-point residuals are displayed below with the same colour code. *Rg* distributions computed for DLD3 (D), DLD10 (E) and DLD17 (F) using the ensembles computed using the different modelling strategies. Equivalent figures for the complete benchmark are provided in S.I. (see Figures S4, S5 and S6).

The analyses of the resulting curves showed that these proteins presented a broad range of sizes, with *R*_*g*_ values spanning from 26.6 Å to 33.4 Å, with a slow but systematic increase with the length of the linker. Interestingly, DLD15, which has a proline-rich 33-residue long linker, exhibited a larger *R*_*g*_ value than DLD18 that has a much longer linker containing 88 residues. Moreover, another protein with a proline-rich linker, DLD10, also presented a larger *R*_*g*_ than other proteins with the same length. Similarly to the *R*_*g*_values, the maximum intramolecular distance, *D*_max_, derived from the pairwise distance distribution, *p*(*r*), showed a gradual increase with the length of the linker, from 103 Å to 144 Å (see values in Table 1). Although is well known that the *D*_max_ values of flexible systems derived from SAXS experiments are not robust structural descriptors [38, 39], some dependence on the composition of the linker was observed. For instance, despite having the same number of residues, DLD10 presented a larger *D*_max_ (133 Å) than DLD8 and DLD9 (119 Å and 117 Å, respectively). This is in line with the larger *R*_*g*_ also observed for DLD10. Altogether, these observations suggested that conformational features of the individual linkers affect the overall size of DLD proteins in solution.

The inspection of the Kratky plots (Fig. S2 in S.I.) indicated that our benchmark set of DLD proteins is highly extended and flexible. A clear peak was identified for all proteins, but its maximum was displaced from the value expected for a globular particle [40]. Moreover, the Kratky profile did not reach a value of 0 at large angles. These observations unambiguously indicate that the benchmark displays extensive flexibility. The *p*(*r*) functions, with the absence of an isolated second peak corresponding to a dumbbell architecture, and smooth decays towards *D*_max_ values, substantiated the presence of high degree of disorder in these proteins (results not shown) [38].

In summary, the model-free analysis of the SAXS profiles demonstrated that SAXS curves measured for this family of proteins represented an excellent dataset to quantitatively validate computational methods adapted to the modelling of flexible multidomain proteins.

### 2.3 Variable performance of ensemble generation methods in reproducing SAXS data

We applied five computational methods for the generation of conformational ensembles of DLD proteins with the aim to evaluate their accuracy using the previously described SAXS dataset. Concretely, MoMA-FReSa [30, 31], CALVADOS3 [27], Mpipi-Recharged [28], bAIes [32] and BioEmu [41], which are all based on distinct principles, were used in this evaluation. MoMA-FReSa implements a stochastic sampling of disordered regions from local structural information extracted from a database of small protein fragments. CALVADOS3 and Mpipi-Recharged generate ensembles from MD simulations considering one-bead-per-residue coarse-grained (CG) potentials. bAIes also relies on MD simulations, but it uses an all-atom model based on a simplified amber99SB-ILDN force field [42] complemented with a bias potential derived from AlphaFold-predicted pairwise residue distance distributions [43]. Finally, BioEmu is a deep-learning-based approach, trained on a large dataset of MD simulations and experimental data, that generates conformational ensembles directly from protein sequences. Note that for a single case, DLD16, we tested two additional deep-learning-based methods, IDPfold2 [44] and IDPForge [45]. However, based on their poor performance (results not shown), we did not compute ensemble models for the other DLD cases. For each of the methods and for each of the proteins, we generated conformational ensembles of similar size (around 10,000 conformations). It should be noted that computational requirements varied considerably depending on the method used. On the hardware employed in this study, MoMA-FReSa required only a few minutes on a multi-core CPU, whereas the coarse-grained MD approaches CALVADOS3 and Mpipi-Recharged typically required on the order of several hours. The all-atom bAIes simulations were substantially more demanding, with runtimes on the order of tens of hours on a single CPU core. BioEmu generated ensembles within a few hours on a modern GPU. Importantly, for MD-based methods, we ran several replicas to verify that the simulation reached convergence with respect to the SAXS data (see Methods section and Figures S3a-c). For the comparative analyses, we used the individual conformational ensemble that provided the best fit to SAXS profile.

Averaged simulated SAXS profiles were computed from each of the ensembles using Crysol [46] and directly compared with the experimental curves using the reduced *χ*^2^-value to quantitatively evaluate the capacity of the different modelling approaches to capture the behaviour of DLD proteins in solution. The performance of the distinct approaches exhibited an enormous disparity (Table 2). Despite its simplicity, MoMA-FReSa was the computational method that showed the best capacity to reproduce the experimental SAXS profiles. Indeed, MoMA-FReSa was the most accurate method in 13 out of the 18 DLD proteins of the benchmark, while CALVADOS3 or Mpipi was the best method for three and two cases, respectively. For MoMA-FReSA, the *χ*^2^-values ranged from 1.87 to 20.15, while this range varied from 3.59 to 42.56 for CALVADOS3, and from 1.98 to 98.47 for Mpipi. Conversely, the other two approaches, bAIes and BioEmu, exhibited a poor capacity to describe the benchmark, with several *χ*^2^-values above 100.

**Table 2:**
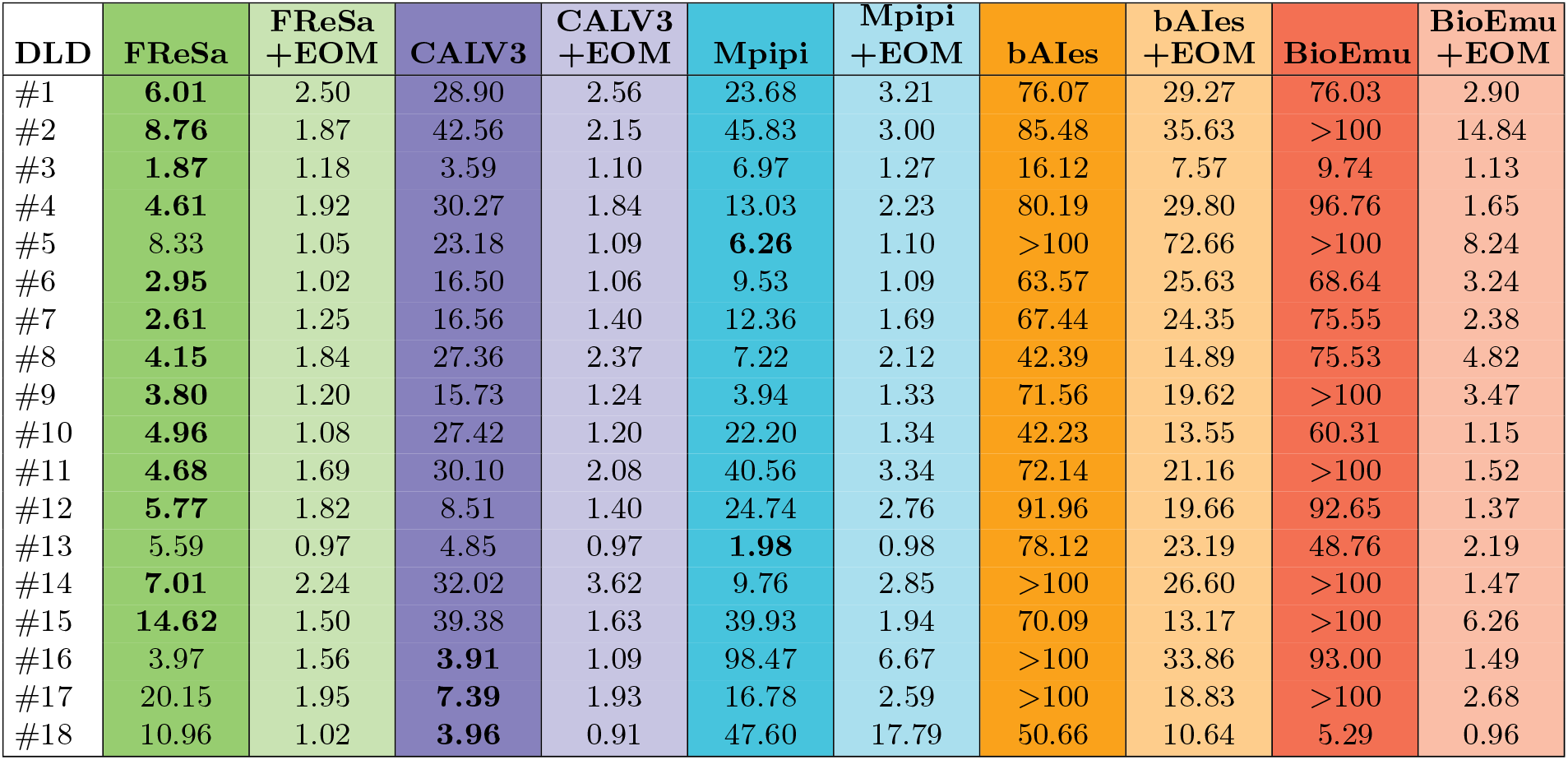
For each of the five ensemble generation methods tested, this table provides the *χ*^2^-values from the comparison of simulated and experimental SAXS profiles. *χ*^2^ values are provided for the ensembles before and after refinement using the EOM method. The best results for each protein before refinement are indicted in bold.

| DLD | FReSa | FReSa<br>+EOM | CALV3 | CALV3<br>+EOM | Mpipi | Mpipi<br>+EOM | bAIes | bAIes<br>+EOM | BioEmu | BioEmu<br>+EOM |
| --- | --- | --- | --- | --- | --- | --- | --- | --- | --- | --- |
| #1 | **6.01** | 2.50 | 28.90 | 2.56 | 23.68 | 3.21 | 76.07 | 29.27 | 76.03 | 2.90 |
| #2 | **8.76** | 1.87 | 42.56 | 2.15 | 45.83 | 3.00 | 85.48 | 35.63 | >100 | 14.84 |
| #3 | **1.87** | 1.18 | 3.59 | 1.10 | 6.97 | 1.27 | 16.12 | 7.57 | 9.74 | 1.13 |
| #4 | **4.61** | 1.92 | 30.27 | 1.84 | 13.03 | 2.23 | 80.19 | 29.80 | 96.76 | 1.65 |
| #5 | 8.33 | 1.05 | 23.18 | 1.09 | **6.26** | 1.10 | >100 | 72.66 | >100 | 8.24 |
| #6 | **2.95** | 1.02 | 16.50 | 1.06 | 9.53 | 1.09 | 63.57 | 25.63 | 68.64 | 3.24 |
| #7 | **2.61** | 1.25 | 16.56 | 1.40 | 12.36 | 1.69 | 67.44 | 24.35 | 75.55 | 2.38 |
| #8 | **4.15** | 1.84 | 27.36 | 2.37 | 7.22 | 2.12 | 42.39 | 14.89 | 75.53 | 4.82 |
| #9 | **3.80** | 1.20 | 15.73 | 1.24 | 3.94 | 1.33 | 71.56 | 19.62 | >100 | 3.47 |
| #10 | **4.96** | 1.08 | 27.42 | 1.20 | 22.20 | 1.34 | 42.23 | 13.55 | 60.31 | 1.15 |
| #11 | **4.68** | 1.69 | 30.10 | 2.08 | 40.56 | 3.34 | 72.14 | 21.16 | >100 | 1.52 |
| #12 | **5.77** | 1.82 | 8.51 | 1.40 | 24.74 | 2.76 | 91.96 | 19.66 | 92.65 | 1.37 |
| #13 | 5.59 | 0.97 | 4.85 | 0.97 | **1.98** | 0.98 | 78.12 | 23.19 | 48.76 | 2.19 |
| #14 | **7.01** | 2.24 | 32.02 | 3.62 | 9.76 | 2.85 | >100 | 26.60 | >100 | 1.47 |
| #15 | **14.62** | 1.50 | 39.38 | 1.63 | 39.93 | 1.94 | 70.09 | 13.17 | >100 | 6.26 |
| #16 | 3.97 | 1.56 | **3.91** | 1.09 | 98.47 | 6.67 | >100 | 33.86 | 93.00 | 1.49 |
| #17 | 20.15 | 1.95 | **7.39** | 1.93 | 16.78 | 2.59 | >100 | 18.83 | >100 | 2.68 |
| #18 | 10.96 | 1.02 | **3.96** | 0.91 | 47.60 | 17.79 | 50.66 | 10.64 | 5.29 | 0.96 |

The excellent description of the structural behaviour of DLD proteins provided by the MoMA-FReSa ensembles indicated that the selected linkers present a high level of disorder. Note that this computational approach samples the conformational space in a sequence-dependent manner, but neither the electrostatic nor the hydrophobic long-range interactions are considered when building ensembles. Interestingly, DLD17, the protein for which MoMA-FReSa performed the worst, with *χ*^2^ = 20.15, compared with CALVADOS3, with *χ*^2^ = 7.39, encompassed the linker with the highest overall net charge, -9 (Table 1). We hypothesized that the linker experienced electrostatic interactions with the globular domains that were captured by CALVADOS3, but not by MoMA-FReSa. Similarly, CALVADOS3 performed notably better than MoMA-FReSa for DLD18, which has the longest linker of the benchmark. Probably, the long linker of this DLD enabled more freedom to establish interactions between both globular domains than the other constructs of the benchmark, and these were better captured by a MD-based approach such as CALVADOS3. Although it is also a CG-MD method, Mpipi did not accurately capture these intramolecular interactions.

The comparison of the synthetic SAXS curves derived from the ensembles with the experimental data provided some clues for the disparity of results obtained for the five modelling approaches (Figure 2A-C, S5 and S6). While bAIes yielded profiles corresponding more extended particles than those present in solution, BioEmu yielded opposite results, generating very compact structures. Interestingly, these features occurred for all the cases of our benchmark, regardless of the composition and length of the linker. Less systematic results were observed for MoMA-FReSa, CALVADOS3, and Mpipi, which produced conformations that were more or less compact than those derived from SAXS measurements, depending on the linker sequence.

To structurally evaluate the ensembles derived from the five methods, we computed their *Rg* distributions (Fig. 2D-F and S6). This analysis indicated strong conformational bias depending on the computational approach. On the one hand, for all DLD proteins, BioEmu exhibited a strong enrichment in highly compact conformations, with a highly populated peak corresponding to conformations with small *Rg* values. However, extended conformations were also populated, although less frequently. Mpipi displayed a similar behaviour for DLD16 and DLD18, showing a strong bias towards highly compact structures. On the other hand, bAIes built ensembles sampling a broad range of sizes, with an enrichment in extended conformations, while compact structures were scarcely sampled. MoMA-FReSa, CALVADOS3 and Mpipi built more balanced ensembles sampling a broad range of protein sizes. Interestingly, CALVADOS3 and Mpipi ensembles systematically showed an enrichment in more compact species than MoMA-FReSA, which did not show any special conformational bias, as expected for this type of stochastic sampling method. Similar features were observed when monitoring the inter-domain distances computed between the centers of mass (CoM) of the individual domains (Figure S7). Note that BioEmu for all cases and Mpipi for some of them overpopulated conformations where the two domains were specifically interacting. Conversely, conformations built with bAIes did not show contacts between globular domains.

Altogether, these analyses showed that modelling tools displaying a balanced population between compact and extended conformations provided the best accuracy to the DLD SAXS dataset. Conversely, these methodologies presenting systematic structural biases resulted in inaccurate ensembles that were unable to reproduce the behaviour in solution of our protein benchmark set.

### 2.4 Can SAXS-guided refinement rescue structurally biased ensembles?

Conformational ensembles are often used as starting pools for data-driven refinement using distinct computational approaches [47]. Here, we have used the conformational ensembles generated by the five computational approaches as starting pools to fit the previously described experimental SAXS curves with the GAJOE algorithm of the Ensemble Optimization Method (EOM) [48, 49], the most popular approach to describe SAXS data of flexible systems. With this analysis, we aimed at exploring the capacity of experimental SAXS data to compensate for large structural disparities observed among the ensembles. The same EOM procedure, searching for the optimal 50-structure sub-ensemble, was applied in all cases (see Methods for details).

The results of this analysis, summarized in Table 2, revealed two distinct trends. High-quality ensembles, *i*.*e*. with low *χ*^2^-values, were obtained from the pools generated by MoMA-FReSa, CALVADOS3, Mpipi and, in some cases, by BioEmu. Concretely, from MoMA-FReSa, CALVADOS3 and Mpipi initial pools, sub-ensembles describing the SAXS curves with good to excellent *χ*^2^-values could be derived for almost all proteins of the benchmark set. Examples of these high quality EOM fits are displayed in Figures 3, S8 and S9. Importantly, when ensembles could be re-weighted to properly describe the data (*χ*^2^-values < 4.0), the capacity of the four building approaches to describe the SAXS curves was similar, regardless to their initial distance to the behaviour of the protein in solution. In contrast, EOM could not find ensembles with the capacity to accurately describe the SAXS data from initial pools built with bAIes, for which no *χ*^2^-values below 7.6 could be derived with EOM.

**Figure 3:**
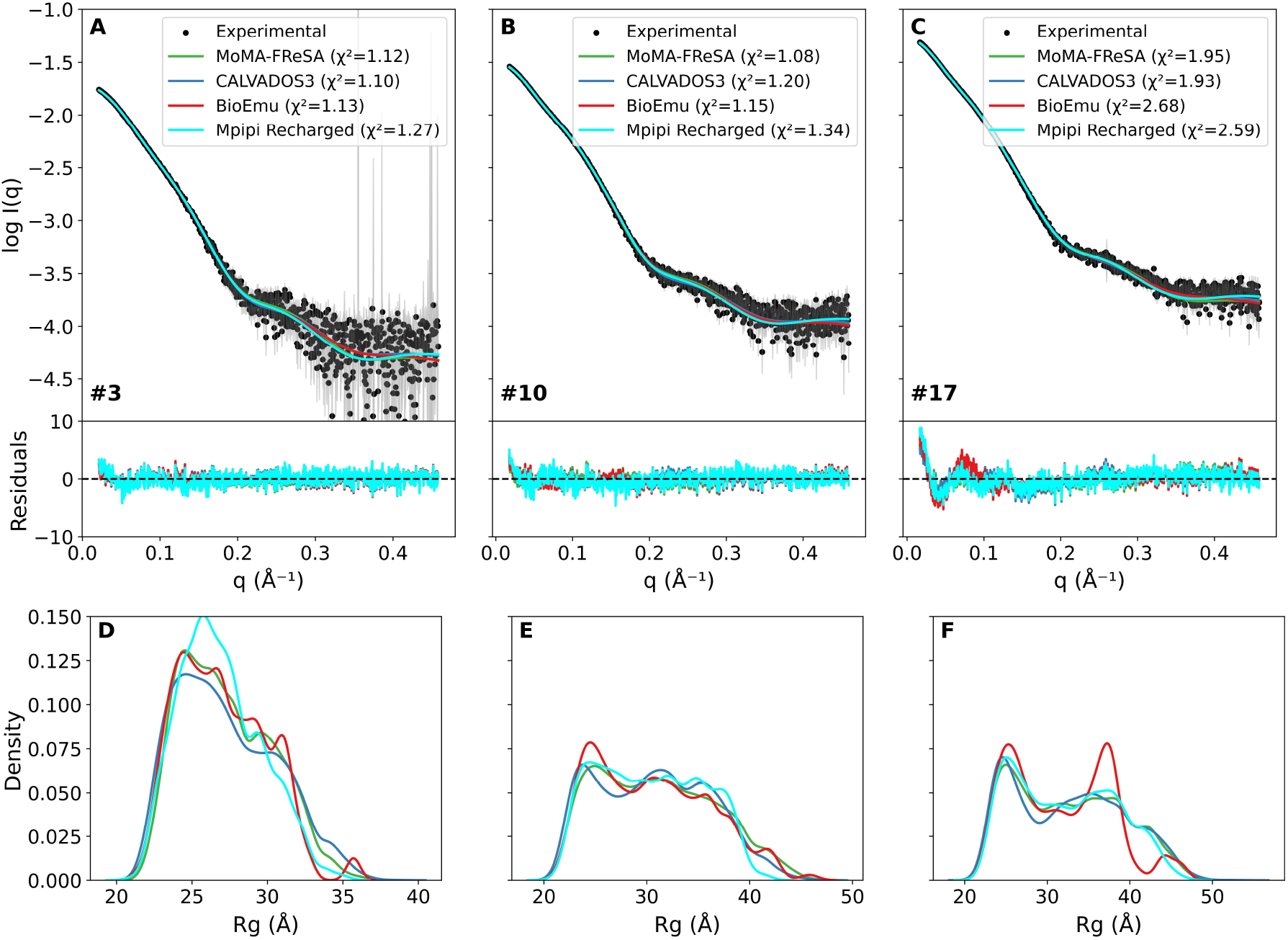
SAXS intensity profiles (black) for DLD3 (A), DLD10 (B) and DLD17 (C) compared with the EOM-fitted scattering profiles computed from the ensembles modelled with MoMA-FReSa (green), CALVADOS3 (blue), Mpipi (cyan) and BioEmu (red). *χ*^2^-values are shown in each panel. Point-by-point residuals are displayed below with the same colour code. *R*_*g*_ distributions for DLD3 (D), DLD10 (E) and DLD17 (F) using the ensembles derived with EOM using the different modelling strategies. Equivalent figures for the complete benchmark are provided in the SI (see Figures S8-S10).

In the previous section, we have shown that BioEmu and bAIes built the most structurally biased ensembles of our benchmark, featuring highly compact and extended conformations, respectively. In contrast, MoMA-FReSa CALVADOS3 and Mpipi provided the most balanced distribution of conformations in terms of *Rg* and distance between both domains. These observations unambiguously demonstrated that the properties of the original ensemble strongly affect the capacity of ensemble-refining programs to find an appropriate subgroup of conformations describing SAXS data. To have a more detailed picture of the effect of the structural biases of the initial pools to fit SAXS data, we analysed the *Rg* distribution of the EOM-fitted ensemble for the DLD proteins (Figure 3, SF). Note that only cases in which EOM was able to correctly fit the data (*χ*^2^ < 4.0) have been plotted [15, 20].

The analysis of the *Rg* distributions indicated that, even when the initial ensemble distributions did not fully agree with the SAXS measurements, a subset of conformations within the pool may still provide a good fit to the experimental profiles. This behavior was particularly evident for CALVADOS3, Mpipi and BioEmu-generated ensembles across many of the DLD proteins in our benchmark set. For these methods, the experimental data were able to correct the original structural bias. Conversely, in all cases for bAIes and in two cases for Mpipi (DLD16 and DLD18), the generated ensembles did not adequately sample all the regions of the conformational space explored by the DLD proteins in solution (Figure S6). Indeed, the significantly reduced exploration of a region of the conformational space could not be compensated by the application of SAXS data. These observations highlighted the need of an exhaustive exploration of the structural landscape in order to obtain a satisfactory description of experimental data.

### 2.5 Do different methods structurally converge after SAXS refinement?

Finally, we studied the similarity between SAXS-refined sub-ensembles derived from distinct original pools. This analysis provided clues on the capacity of SAXS data in order to robustly define the conformational behaviour of proteins in solution. To gain a more intuitive understanding of the conformational differences between ensembles, we examined their corresponding *R*_*g*_ distributions. Figure 3D-F shows the results for three representative cases, DLD3, DLD10 and DLD17, while the distributions for the complete benchmark set are shown in Figure S10. Interestingly, the *R*_*g*_ distributions after the SAXS refinement were very similar, regardless of the conformational features of the original pool. This was particularly true for MoMA-FReSa, CALVADOS3 and Mpipi, which presented astonishingly similar final *R*_*g*_ distributions with very similar population maxima. Although BioEmu also provided distributions globally similar to those of the three methods after EOM-based refinement, they tended to be more spiky and presented discontinuities, especially for very extended conformations. This can be observed for DLD17 (Figure 3F), where three *R*_*g*_ peaks were identified and the last one, at *R*_*g*_ around 47 Å, was detached of the rest of the distribution. We attributed the differences in the *R*_*g*_ distribution derived from BioEmu ensembles to the larger structural distance between the original pool and that sampled by the protein in solution, as we have shown in the previous section (Table 2), and the lower density of conformations with large *R*_*g*_ s.

The similarity of the *R*_*g*_ distribution shapes and the position of the highly populated *R*_*g*_ values of the three refined ensembles suggested that the relative domain positions of the selected poses were consistent across the protein building strategies. In order to have a detailed perspective of this hypothesis, we analysed the CoM distribution for the SAXS-refined ensembles (Figure S11). Compared with the original distributions (Figure S7), the overall shapes of the CoM profiles derived from the four pools were very similar, suggesting that SAXS data were effective in selecting similar structures. When analysing these distributions in more detail, we observed that proteins with short linkers (< 22 residues, from DLD1 to DLD7) exhibited larger disparities among the CoM distributions than those with large linkers. From DLD8, the CoM distributions were very similar, with MoMA-FReSa, CALVADOS3 and Mpipi yielding virtually equivalent profiles for most of the cases. More disparities were observed for BioEmu CoM distributions. Again, we attributed the differences observed in BioEmu to the strong structural differences of its original ensemble with respect to MoMA-FReSa, CALVADOS3 and Mpipi. All together, our analysis of the *R*_*g*_ and CoM distributions suggested that conformational biases of the original conformational pools slightly affect the structural features of the SAXS-refined ensemble.

## 3 Discussion

The present study underscores the substantial variability that still characterizes current approaches for modeling ensembles of disordered proteins [26]. Across the family of DLD proteins analyzed, performance, quantified through comparison with high-quality SAXS data, was strongly dependent on both the computational method employed and the specific linker sequence. The modeling tools tested, which represent distinct families of ensemble-generation approaches, systematically exhibited method-specific structural biases. In particular, BioEmu produced highly compact ensembles, whereas bAIes consistently yielded overly extended conformations. In contrast, ensembles generated using MoMA-FReSA, CALVADOS3 and Mpipi were more balanced and less affected by pronounced structural biases. Among the approaches evaluated, MoMA-FReSA provided the most reliable overall performance, while also offering a very favorable balance between accuracy and computational efficiency. Importantly, the design of our protein dataset enabled us to relate method performance to specific structural features. Both linker composition and linker length emerged as critical determinants of accuracy in a clearly method-dependent manner. For example, CALVADOS3 systematically produced poor fits to SAXS data (large *χ*^2^-values) for proteins encompassing proline-rich (DLD2, DLD10, and DLD15) or glycine-rich (DLD1, DLD8, and DLD14) linkers. This likely reflects limitations of the one-bead-per-residue representation in CALVADOS3, which appears insufficient to capture the distinctive conformational behaviour of these amino acids. In contrast, MoMA-FReSA, which is based on amino-acid-specific conformational sampling derived from high-resolution experimental structures [31], is less sensitive to linker sequence composition. Linker length also significantly influenced method performance. For the six longest linkers analyzed (33 residues or more), CALVADOS3 outperformed the other methods in three cases (Table 2). We speculate that longer linkers may facilitate transient interdomain interactions, which are more readily captured by the CALVADOS3 force-field, resulting in relatively compact ensembles. Interestingly, Mpipi, which is also a CG-MD method, did not properly reproduce data measured for the longest DLD proteins, suggesting that its force-field is less adapted to our benchmark dataset.

The dataset used to benchmark the computational methods spans a broad range of linker sequences and lengths, ensuring a representative diversity of systems. In addition, the presence of two distinct globular domains at the termini breaks the intrinsic symmetry of the proteins, enabling a more rigorous evaluation of the resulting ensemble models. However, this specific DLD architecture may introduce method-dependent biases. In particular, approaches that rely on predicted residue-residue distance distributions may systematically underperform when domain-domain or linker-domain interactions are inaccurately predicted, as it is likely the case for bAIes. Despite these inherent limitations of the benchmark set, our results clearly indicate that no single method consistently reproduces SAXS data across the full set of proteins analyzed. This observation highlights the persistent challenges associated with predicting conformational ensembles for flexible, multidomain systems.

The use of SAXS-guided refinement further illustrates both the strengths and limitations of integrating experimental data with computational ensemble generation. In all cases, refinement resulted in a clear improvement in the agreement with experimental scattering profiles, demonstrating that SAXS data can effectively steer conformational ensembles toward more realistic distributions. However, the quality of the refined ensembles was strongly dependent on the generative method used to construct the initial conformational pool. Methods that produced physically plausible and structurally diverse starting ensembles, most notably MoMA-FReSA, CALVADOS3 and Mpipi, benefited the most from refinement, yielding highly accurate ensembles with low *χ*^2^-values. Interestingly, in some cases BioEmu ensembles, despite being initially far from the solution behavior of the proteins, were able to achieve good agreement with the SAXS data after refinement. This behavior was not observed for bAIes. We attribute the relative success of BioEmu to its broad conformational sampling, which includes compact structures and, to a lesser extent, more expanded conformations (Figures 2D–F and S6). Such diversity appears to provide sufficient flexibility for refinement to effectively reweigh the ensemble toward experimentally consistent states. Overall, our results indicate that a broad and representative sampling of the conformational landscape of disordered proteins is a prerequisite for effective SAXS refinement strategies. The conclusions presented here were obtained using EOM, a maximum-parsimony-based approach. Similar dependencies on the quality of the prior ensemble are expected for maximum-entropy-based refinement methods [50, 51], in which a large fraction of conformations of the initial ensemble are reweighted to achieve agreement with experimental data.

The comparison of SAXS-refined ensembles with low *χ*^2^-values derived from different initial conformational pools provided further insight into both the robustness and the limitations of current modeling and analysis strategies. Notably, ensembles generated using distinct modeling approaches converged upon SAXS refinement. Specifically, SAXS-refined ensembles originating from MoMA-FReSa, CALVADOS3 and Mpipi displayed remarkably similar *R*_*g*_ and CoM distributions, not only preserving the overall range of sampled radii of gyration but also reproducing comparable fine structural features (Figures 3D–F and S10). Strikingly, even BioEmu, whose original conformational pool differed substantially from those of MoMA-FReSa, CALVADOS3 and Mpipi, yielded SAXS-refined ensembles with closely matching *R*_*g*_ distributions and similarly populated states. Together, these observations underscore the strong constraining power of SAXS data in defining robust distributions of global structural parameters for highly flexible proteins. However, the intrinsically low-resolution nature of SAXS data necessitates cautious interpretation of refined ensembles. While SAXS can reliably constrain overall molecular dimensions and interdomain CoM distances, it does not provide sufficient information to uniquely determine specific interdomain interactions or detailed local structural features [15, 20]. Yet, our results underline the power of SAXS data to drive computational approaches to identify residue-specific contacts and provide a high-resolution picture of stable conformers present in solution.

Beyond methodological considerations, the biological and biotechnological relevance of accurate ensemble modeling deserves particular emphasis. Flexible linkers play a central role in mediating the function of modular proteins by regulating domain spacing, orientation, and dynamics [1, 6]. Reliable ensemble predictions are therefore essential for understanding how these proteins operate in their native biological contexts. Furthermore, prediction accuracy also has direct implications for the engineering of multimodular enzymes for biotechnological applications, where linker design and domain organization can be exploited to optimize catalytic activity, specificity or stability. As demonstrated in this study, their functional and biotechnological importance contrasts sharply with the persistent difficulties in accurately predicting their structural and dynamical properties. In this regard, the integration of experimental data with physics-based modeling approaches emerges as a particularly promising strategy. Crucially, predicted ensembles should be systematically validated against experimental data that probe distinct and complementary structural features. Such validation is necessary to ensure that the resulting ensembles faithfully capture both the underlying energetic landscape and the conformational preferences encoded in protein sequences.

## 4 Materials and Methods

### Selection of linkers from natural diversity

Sequences of multimodular enzymes containing catalytic GH11 and CBM domains were retrieved from the CAZy database [34]. To ensure consistency, we selected only linkers flanked by a GH11 domain at the N-terminus and a CBM at the C-terminus. Sequences originating from eukaryotes and bacteria were retained, and an additional length constraint was imposed by selecting linkers between 10 and 100 amino acids. After redundancy filtering, this procedure yielded a total of 769 linker sequences.

For each linker, physicochemical properties were computed using localCIDER [35]. The resulting feature set was used to cluster the sequences via hierarchical agglomerative clustering (HAC) with Ward’s linkage method [52]. Based on the resulting dendrogram, we selected 18 representative linker sequences that spanned the observed diversity in terms of length and physicochemical composition. This reduced yet diverse subset was chosen to capture the main structural and compositional trends present in natural GH11–CBM linkers, while remaining experimentally and computationally tractable for downstream analyses.

### DNA Constructs

The synthetic genes for the glycosyl hydrolase (GH) catalytic module from *Neocallimastix patriciarum*, and for a fusion protein comprising the DLD16 linker and the carbohydrate-binding module (CBM) from *Cellulomonas fimi* were purchased at IDT® (Integrated DNA Technologies). Linker sequences corresponding to DLD1 to DLD17 (excluding DLD16) were ordered as Ultramer® Duplex DNA fragments at IDT®. Due to their length and complexity, the linker corresponding to DLD18 was purchased at GeneArt® (Life Technologies®).The GH gene was first cloned in-frame into the pDB-6His-ccdb plasmid using NcoI and NdeI restriction sites, generating the construct pDB-6His-GH-ccdb. The DLD16–CBM fusion was then inserted between the NdeI and BlpI sites to obtain the pDB-6His-GH-DLD16-CBM plasmid. The insertion and exchange of the different linker sequences (DLD1–18) were performed by an In-Fusion reaction between the NdeI and XhoI restriction sites located between the GH and CBM coding regions.

### Protein production and purification

The pDB-6His-GH-DLDXX-CBM vectors were transformed in *E. coli* BL21(DE3) strain and grown in LB medium supplemented with 50 µg/mL of kanamycin until the OD reached 0.6. Protein expression was induced by the incorporation of IPTG (0.5 mM), and cells were grown for 3 h before harvesting them by centrifugation for 20 min at 6000g at 4°C. Pellets were resuspended in Tris-HCl 20 mM pH 7.5, NaCl 300 mM and 2 mM DTT (buffer A) and stored at -80°C. Cells were supplemented with a Complete® EDTA free tablet (Roche) and lysed by sonication. Insoluble proteins and cell debris were sedimented by centrifugation at 40000g at 4°C for 30 min. Supernatant was supplemented with imidazole to a 5 mM final concentration and loaded onto a 5 ml affinity column (HisTrap Excel, Cytiva) equilibrated with buffer B (buffer A containing 5 mM imidazole). The column was washed with buffer B and proteins were eluted with a linear 0 to 100% gradient of buffer C (buffer A containing 0.5 M imidazole). The peak fractions were analyzed by SDS-PAGE. Fractions containing tagged GH-linker-CBM fusion protein were pooled and dialyzed overnight at 4 °C against buffer D (Na2HPO4/NaHPO4 50 mM pH 6, NaCl 50 mM, DTT 2 mM). The dialyzed protein was then loaded on a Superdex S75 16/60 (HiLoad 16/600 Superdex 75 pg, Cytiva) equilibrated with buffer E (Na2HPO4/NaHPO4 50 mM pH 6, TCEP 2 mM). Fractions containing the protein were analysed by SDS-PAGE and pooled. The purified proteins were concentrated to approximately 10 mg/mL with Vivaspin centrifuge concentrator (Sartorius stedim biotech) before structural analyses.

### Small-Angle X-ray Scattering measurements

Small-Angle X-ray Scattering (SAXS) experiments were performed on the purified DLD constructs at the SWING beamline of the SOLEIL Synchrotron (Saint-Aubin, France) [53], using an X-ray wavelength of 1.03 Å and a sample-to-detector distance of 2.00 m. Measurements were conducted at 20 °C using Size-Exclusion Chromatography coupled to SAXS (SEC-SAXS) in buffer E. A volume of 45 µL of sample was injected into a 3 mL Superdex 200 5/150 GL column (Cytiva, Uppsala, Sweden) pre-equilibrated with the same buffer and eluted at a flow rate of 0.2 mL/min. The acquired scattering data covered a momentum transfer range of 0.002 Å^*−*1^ < q < 0.5 Å^*−*1^. Buffer scattering was recorded before the void volume of the column (1 mL). Data processing was carried out using CHROMIX [54], which automatically identified buffer and sample frames, subtracted the buffer signal, and averaged the scattering profiles. Final SAXS curves were analyzed using Primus from the ATSAS software suite [55].

### Ensemble generation methods

#### Definition of rigid and flexible regions

Several of the ensemble-generation methods applied in this work require prior specification of rigid and flexible regions. Figure S12 illustrates the generic amino acid composition of the DLD constructs, in which the linker sequence varies across the 18 tested constructs. Based on this architecture, the segments MGSSHHHHHH, TTHM–linker–LESTGC, and SGGGTA were defined as flexible, while the remaining regions were treated as rigid. This partitioning of flexible and rigid residues was consistently applied in the MoMA-FReSA, CALVADOS3, and Mpipi-Recharged simulations.

#### MoMA-FReSa

MoMA-FReSA [30,31] requires as input an initial structure for the rigid domains together with a definition of which regions of the sequence are treated as rigid or flexible. Conformational ensembles are generated by sequentially sampling backbone dihedral angles within the flexible regions from distributions observed in naturally occurring coil conformations. We generated ensembles of 10,000 conformations for each of the 18 DLD constructs using the three-residue-based sampling strategy (TRS) strategy, where the identity and structure of the two neighboring residues are also taken into account.

#### CALVADOS3

We performed 1 *µ*s coarse-grained molecular dynamics (CG-MD) simulations for each DLD construct using CALVADOS3 [27] with default parameters. The initial conformations were generated using MoMA-FreSa, with residues belonging to folded domains restrained and the remaining residues treated as flexible (Figure S12). All-atom structures were coarse-grained to a one-bead-per-residue representation using cg2all [56]. By default, CALVADOS3 employs Langevin dynamics, specifically the LangevinMiddleIntegrator implemented in OpenMM, with a time step of 10 fs. To avoid initial equilibration artifacts, the first 100 ns of each trajectory were discarded. Protein configurations were saved every 100 ps, and the resulting CG ensembles were back-mapped to all-atom representations using PULCHRA [57].

#### Mpipi-Recharged

We performed 1 *µ*s CG-MD simulations using the Mpipi-Recharged force-field [28] for each DLD construction. Simulations were carried out using LAAMPS [58] (2 August 2023 version). Each starting conformation was originally generated by MoMA-FReSa, where the residues of the folded domains were kept fixed and the remaining ones were defined as flexible (see Figure S12). The all-atom starting structures were coarse-grained to a one-bead per residue representation using cg2all [56]. Then, the structure files were converted to the LAAMPS input format using in-house scripts, scaling the interaction parameters of the globular residues accordingly [28]. We performed local energy minimization of each starting structure using the steepest descendent algorithm followed by 20000 equilibration steps in NVT regime with a timestep of 1 fs. Finally, we produced a 1 *µ*s of NVT production using an implicit solvent model, already defined in the Mpipi-Recharged parameters (timestep: 10 fs, temperature: 293K, ionic strength: 150 mM). Snapshots were recorded every 100 ps. The generated ensembles were back-mapped to all-atom using PULCHRA [57].

#### bAIes

We obtained 100 ns of bAIes MD-simulations for each DLD construct [32]. In this case, the starting structures were obtained using AlphaFold 2 on ColabFold [59] together with the corresponding distogram for each protein, which is a mandatory input file for bAIes. Note that the AlphaFold 2 predicted structures of the GH11 and the CBM domains were extremely similar to the experimental structures ones, with RMSDs of 0.340 and 0.692 Å, respectively. In all cases, an energy minimization of each starting structure was carried out using the conjugate gradient algorithm followed by 100 ns of NVT production in vacuum (timestep of 1 fs and temperature of 293K). Snapshots were recorded each 10 ps. SAXS profiles were computed for the last 80 ns of the simulation.

#### BioEmu

BioEmu [41] is a parameter-free method. We ran BioEmu to generate ensembles of 10,000 conformations for each construct. Conformations exhibiting steric clashes were automatically filtered by the algorithm; consequently, the final ensemble size slightly varied across DLD systems.

### Convergence assessment of MD-based methods

For CALVADOS3 simulations, we performed three independent replicas for each DLD, whereas five replicas were generated for Mpipi-Recharged. Convergence was assessed using the cumulative *R*_*g*_ as the primary criterion (see Figures S3a–c). The larger number of replicas for Mpipi-Recharged was motivated by the larger variability observed across independent runs compared to CALVADOS3, and was introduced to ensure the reliability of the results. In all cases, simulations reached convergence for every DLD independently of the starting conformation.

For bAIes, we ran 5 independent replicas for the DLD16 and DLD18 constructions and we assessed the their convergence via the cumulative *Rg* (Figures S3a–c). In each case, one of the starting structures was obtained using AlphaFold 2 on ColabFold [59], while the rest of structures were generated using MoMA-FReSa. In both DLD and for all the starting structures, we used the same distogram retrieved from AlphaFold 2. Given the high degree of convergence of these two examples (Figure S3c, panels K and Q), and due to the higher computational cost compared with the other MD-based approaches, we ran a single trajectory for the rest of the DLD constructs using AlphaFold 2 predictions as starting structures.

### Simulation of SAXS profiles

The SAXS profile for each conformation of all ensembles was calculated using Crysol [46], setting the number of spherical harmonics to 35 and using implicit hydrogens. Both before and after SAXS refinement, the *χ*^2^ value between the theoretical and experimental scattering profiles was calculated as follows:

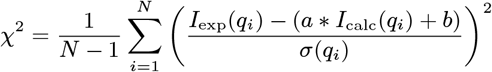

where *N* is the number of experimental data points in the SAXS profile; *I*_exp_(*q*_*i*_) and *I*_calc_(*q*_*i*_) are the experimental and calculated SAXS intensities at momentum transfer *q*_*i*_, respectively; and *σ*(*q*_*i*_) is the experimental error at *q*_*i*_. Finally, *a* and *b* are the scaling factor and constant background offset applied to the theoretical SAXS curve to minimize *χ*^2^.

### Ensemble filtering/refinement (EOM)

The ensembles generated for each DLD were refined using the available SAXS data, using the program GAJOE from the EOM suite [48, 49]. In order to fit the calculated SAXS profiles to experimental data, we ran GAJOE using 200 cycles of 1,000 generations each, with a sub-ensemble size of 50 and disallowing repetitions.

## CRediT authorship contribution statement

**Santiago Rodriguez**: Investigation, Validation, Visualization, Writing – original draft, Writing – review & editing; **Aurélie Fournet**: Investigation, Validation, Writing – review & editing; **Simon Bartels**: Investigation, Writing – review & editing; **Mátyás Pajkos**: Investigation, Software; **Ilinka Clerc**: Software; **Laure Carrière**: Investigation, Software; **Aurélien Thureau**: Investigation, Writing – review & editing; **Cédric Montanier**: Resources, Writing – review & editing, Funding acquisition; **Claire Dumond**: Resources, Writing – review & editing; **Fréderic Allemand**: Supervision, Writing – review & editing; **Juan Cortés**: Conceptualization, Supervision, Writing – original draft, Writing – review & editing, Funding acquisition; **Pau Bernadó**: Conceptualization, Supervision, Writing – original draft, Writing – review & editing, Funding acquisition.

## Data availability

SAXS data will be deposited in the Small Angle Scattering Biological Data Bank (SASBDB; https://www.sasbdb.org) and the SAXS-refined ensembles will be deposited in the Protein Ensemble Database (PED; https://proteinensemble.org). Non-refined ensembles generated by the various methods will also be made publicly available upon publication of the final version of the manuscript.

## Declaration of competing interest

The authors declare that they have no known competing financial interests or personal relationships that could have appeared to influence the work reported in this paper.

## Acknowledgements

We thank Nicolas Terrapon for his assistance with extracting data from the CAZy database. We thank Rosana Collepardo-Guevara (University of Cambridge), Kresten Lindorff-Larsen (University of Copenhagen) and Massimiliano Bonomi (Institut Pasteur) and their group members for useful discussions on the implementation of the computational methods. This work was supported by the French National Research Agency (ANR) under grant ANR-22-CE45-0003 (CORNFLEX project). It was also carried out in the framework of the COST Action ML4NGP [CA21160], supported by the COST (European Cooperation in Science and Technology) and the HORIZON-MSCA-SE project IDPfun2, funded by the European Union under grant agreement no. 101182949. We acknowledge SOLEIL for provision of synchrotron radiation facilities and we would like to thank SWING Staff for assistance in using beamline under BAG proposal 20241146. The CBS is a member of France-BioImaging (FBI) and the French Infrastructure for Integrated Structural Biology (FRISBI), two national infrastructures supported by the French National Research Agency (ANR-10-INBS-04-01 and ANR-10-INBS-05, respectively).

## Notes

### Competing Interest Statement

The authors have declared no competing interest.

### Summary of Updates

The revised version contains new simulations with Mpipi-Recharged in which we have used a more appropriate parametrization. These changes have been performed with the advise of the program developers.

