## Supplementary Information for "Conformational ensembles of flexible multidomain proteins: How close are we to accurate and reliable predictions?"

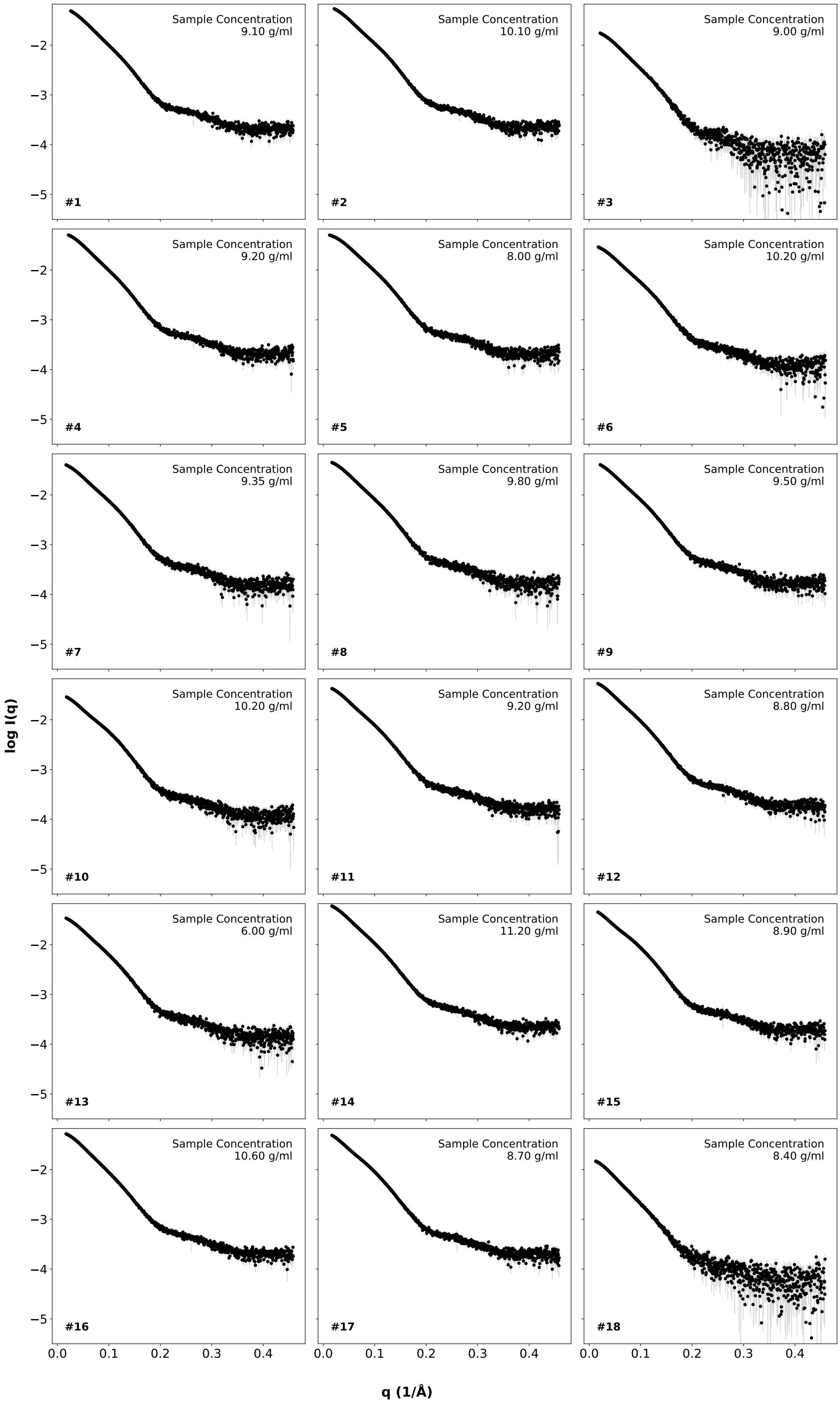


Figure S1. SAXS intensity profiles *versus* the momentum transfer measured for the 18 DLD proteins. The specific DLD number is shown at the bottom left of each panel.

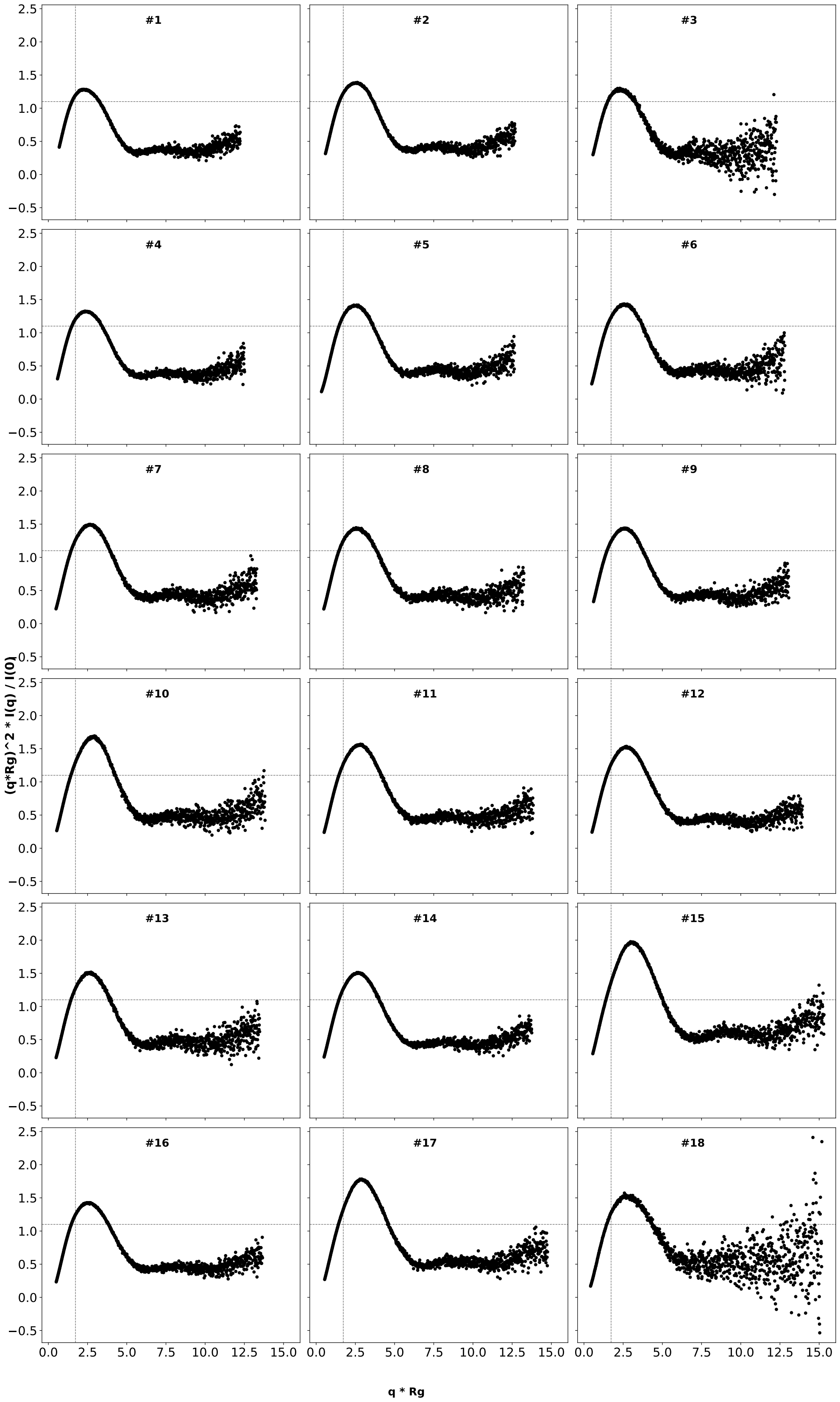


Figure S2. Dimensionless Kratky representation for the 18 DLD proteins measured in this study. The vertical ($\sqrt{3}$) and horizontal (1.104) dashed lines indicate the maximum expected in the Kratky plot for a globular protein. The specific DLD number is shown in each panel.

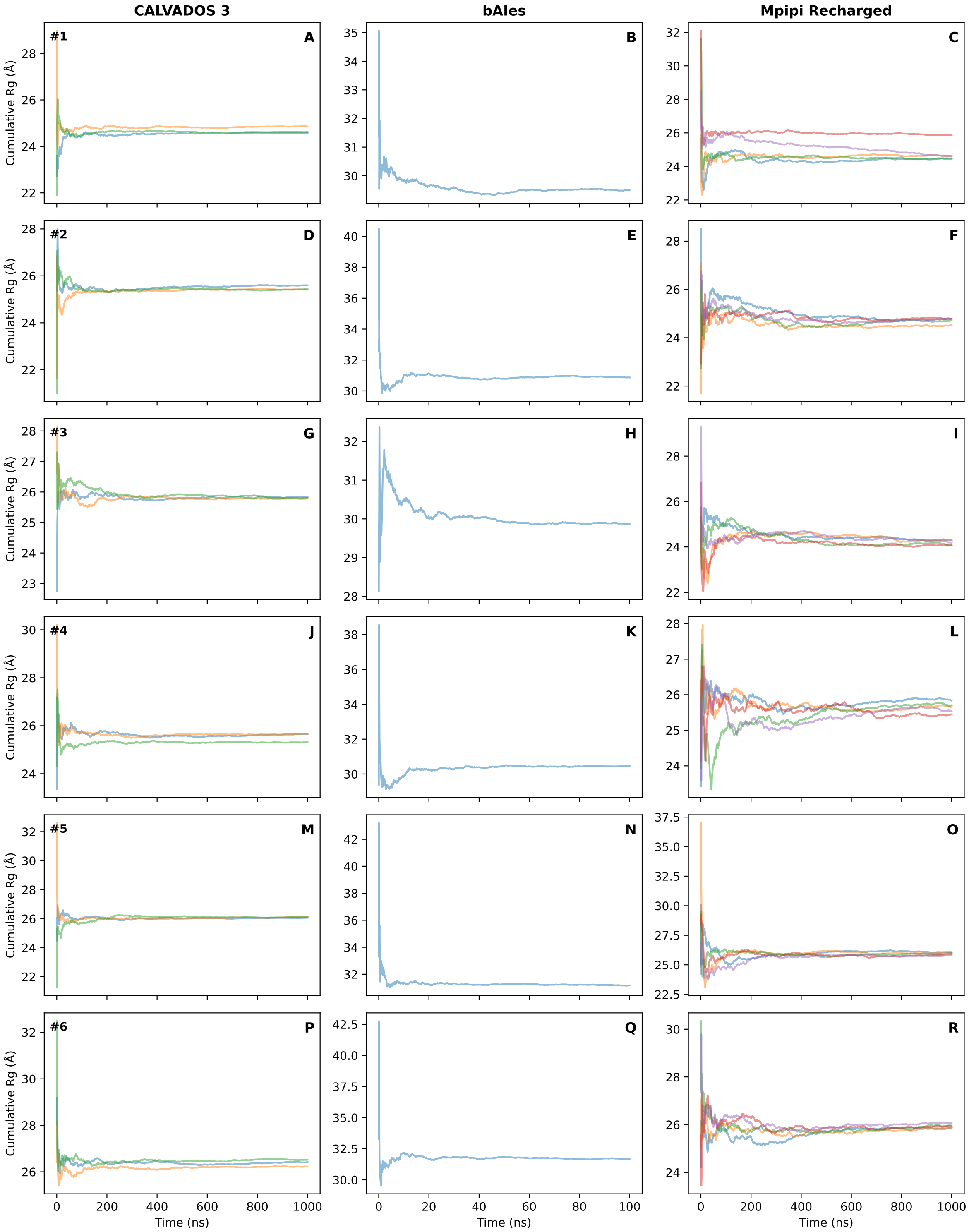


Figure S3a. Cumulative Rg across simulations for the MD-based ensemble generation methods (displayed in the columns). Different colors in each panel represent a different starting conformation in the MD simulation. Note that the scales of the panels change according to the values obtained. The specific DLD number is shown in each row.

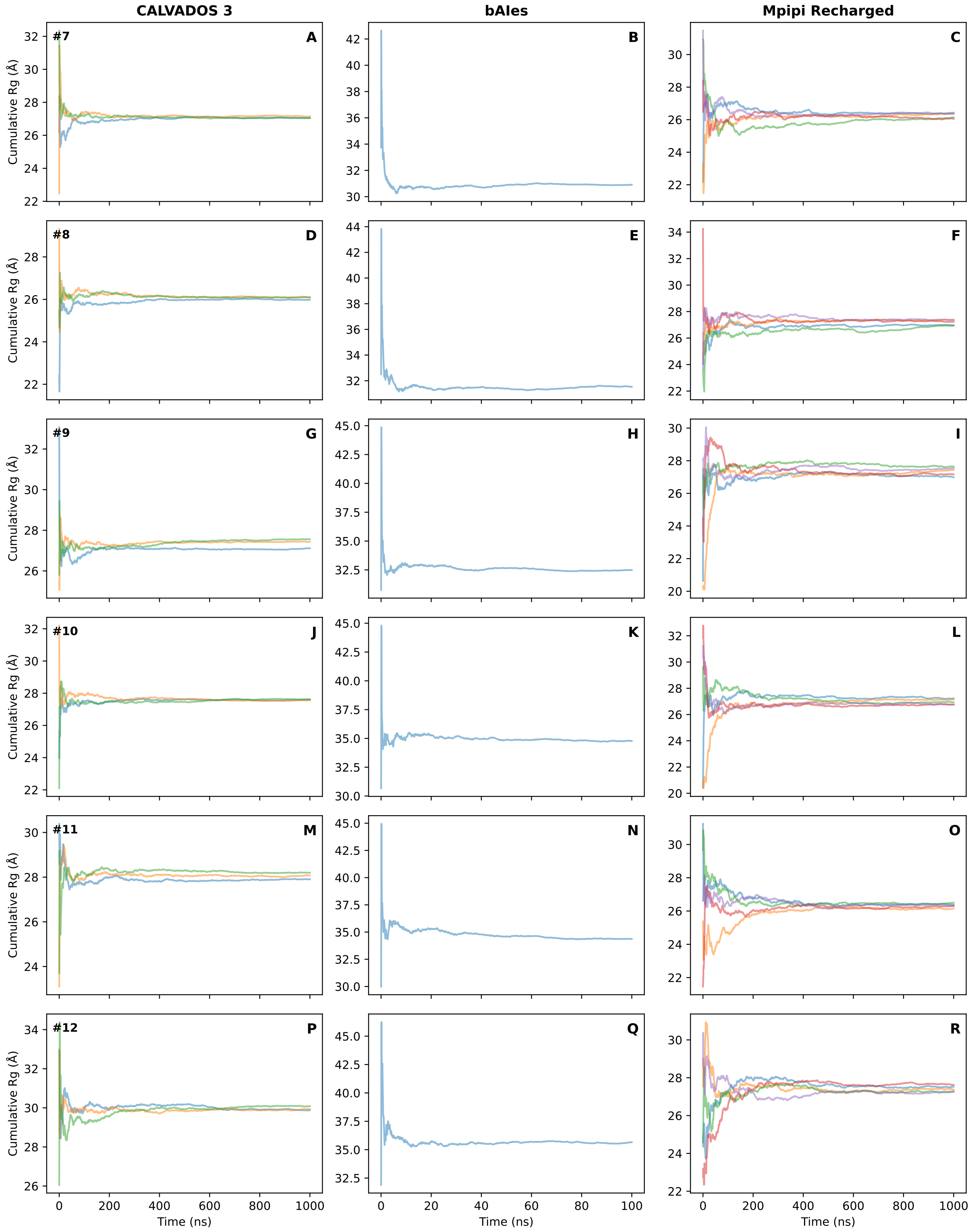


Figure S3b. Cumulative Rg across simulations for the MD-based ensemble generation methods (displayed in the columns). Different colors in each panel represent a different starting conformation in the MD simulation. Note that the scales of the panels change according to the values obtained. The specific DLD number is shown in each row.

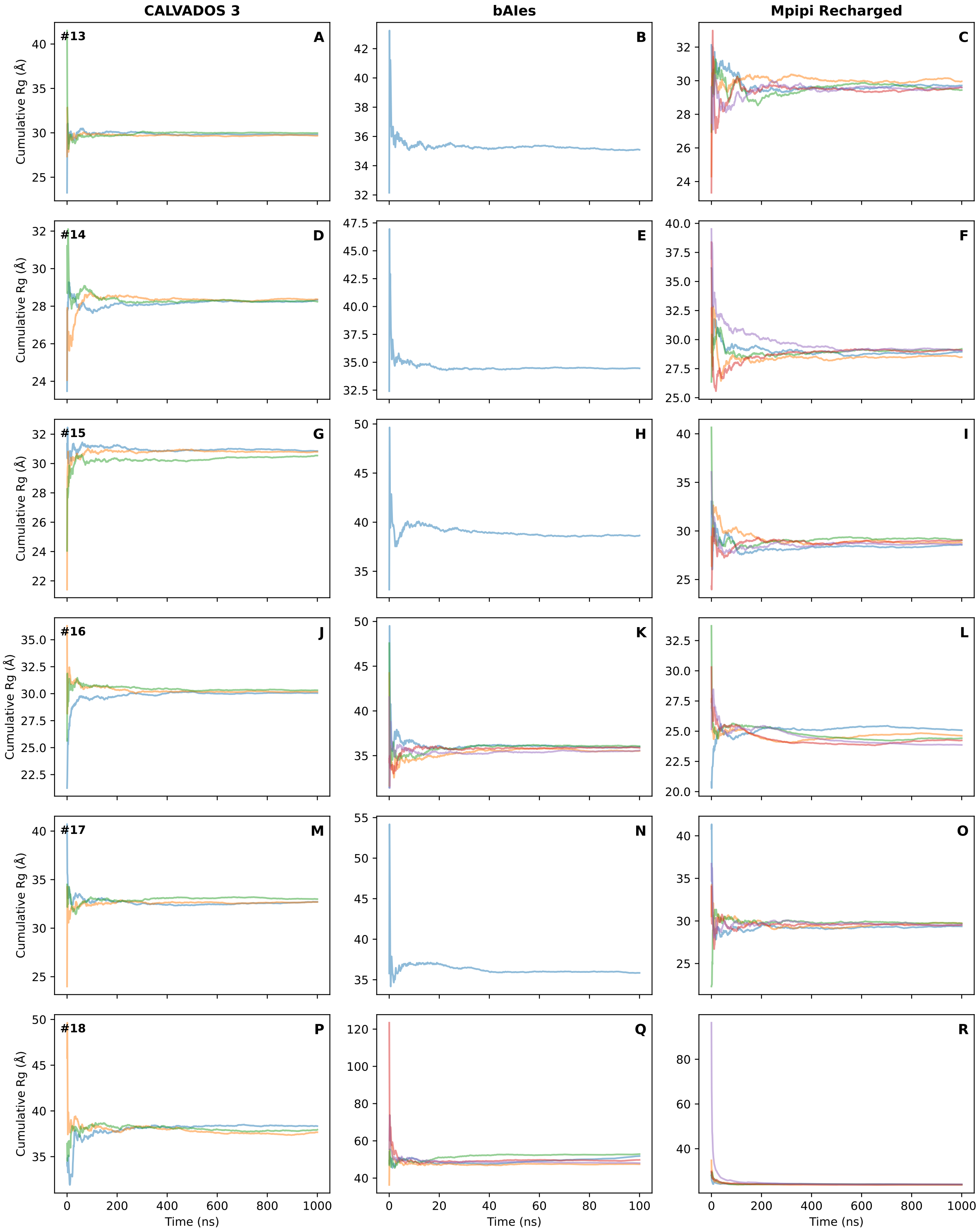


Figure S3c. Cumulative Rg across simulations for the MD-based ensemble generation methods (displayed in the columns). Different colors in each panel represent a different starting conformation in the MD simulation. Note that the scales of the panels change according to the values obtained. The specific DLD number is shown in each row.

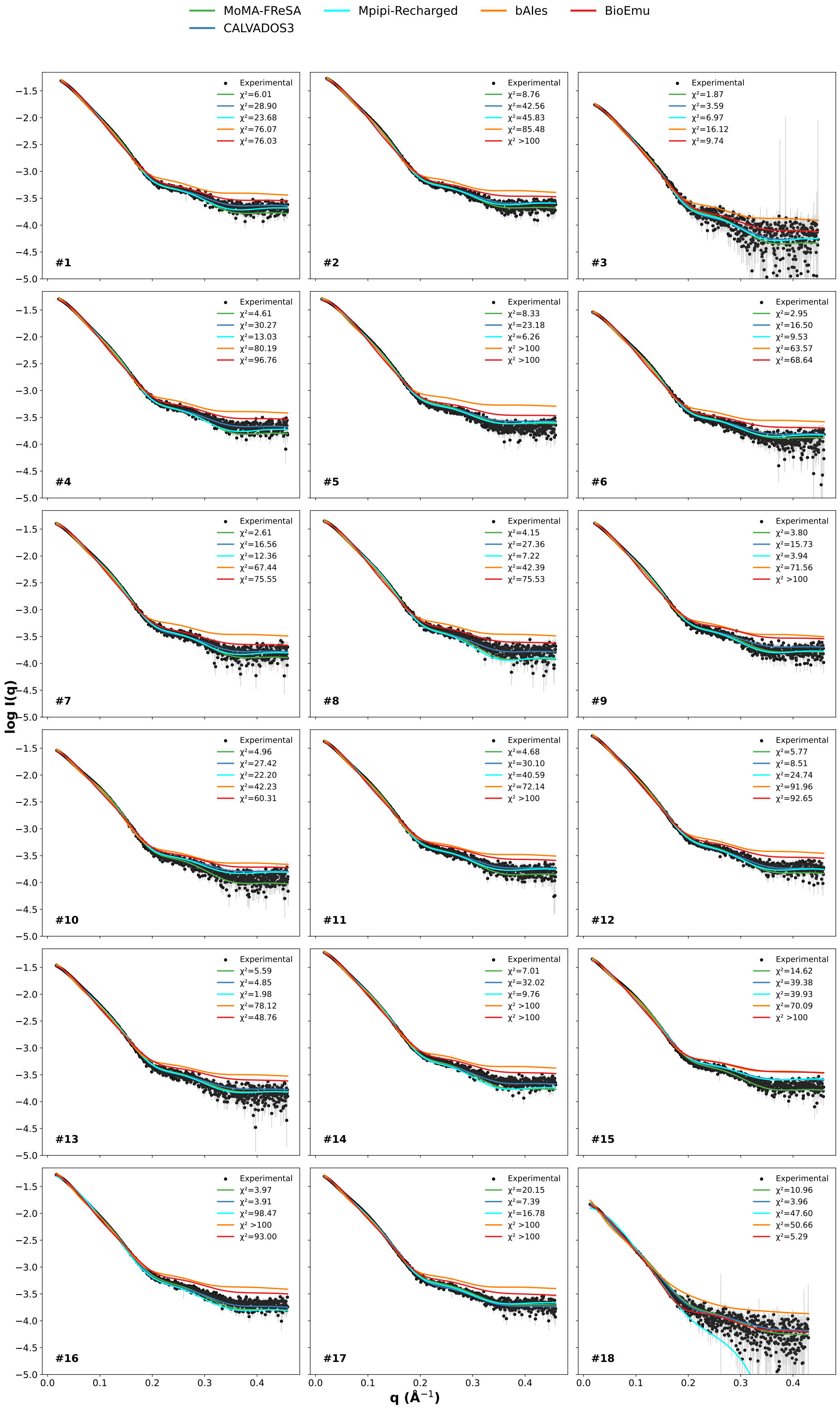


Figure S4. SAXS intensity profiles (black) for all DLDs constructs compared with the averaged scattering profiles computed from the ensembles modelled with MoMA-FReSa (green), CALVADOS3 (blue), Mpipi-Recharged (cyan), bAIes (orange) and BioEmu (red). $\chi^2$-values are shown in each panel. The specific DLD number is shown in each panel.

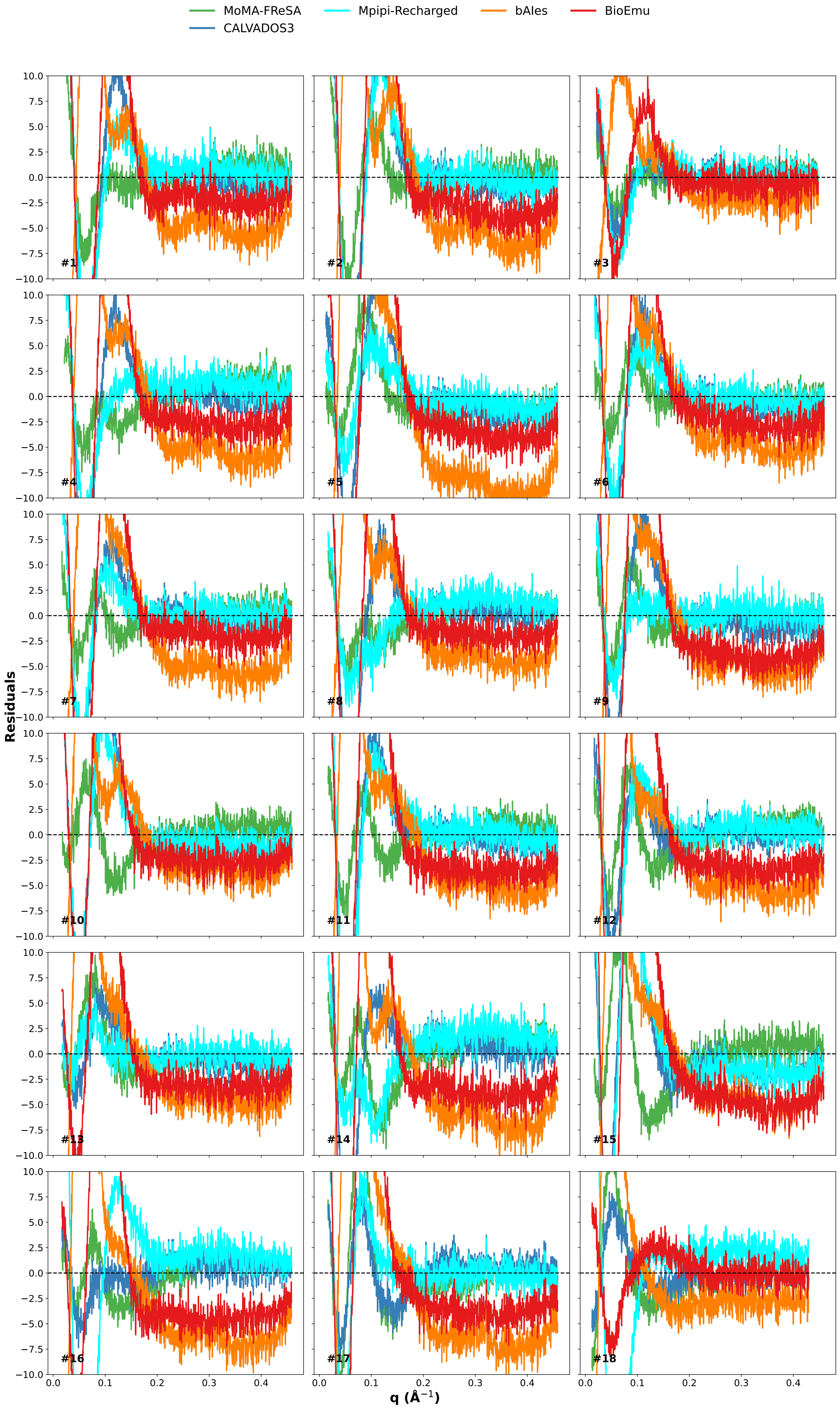


Figure S5. Point-by-point residuals between the SAXS intensity profiles and the averaged theoretical scattering profiles from Figure S4. The specific DLD number is shown in each panel.

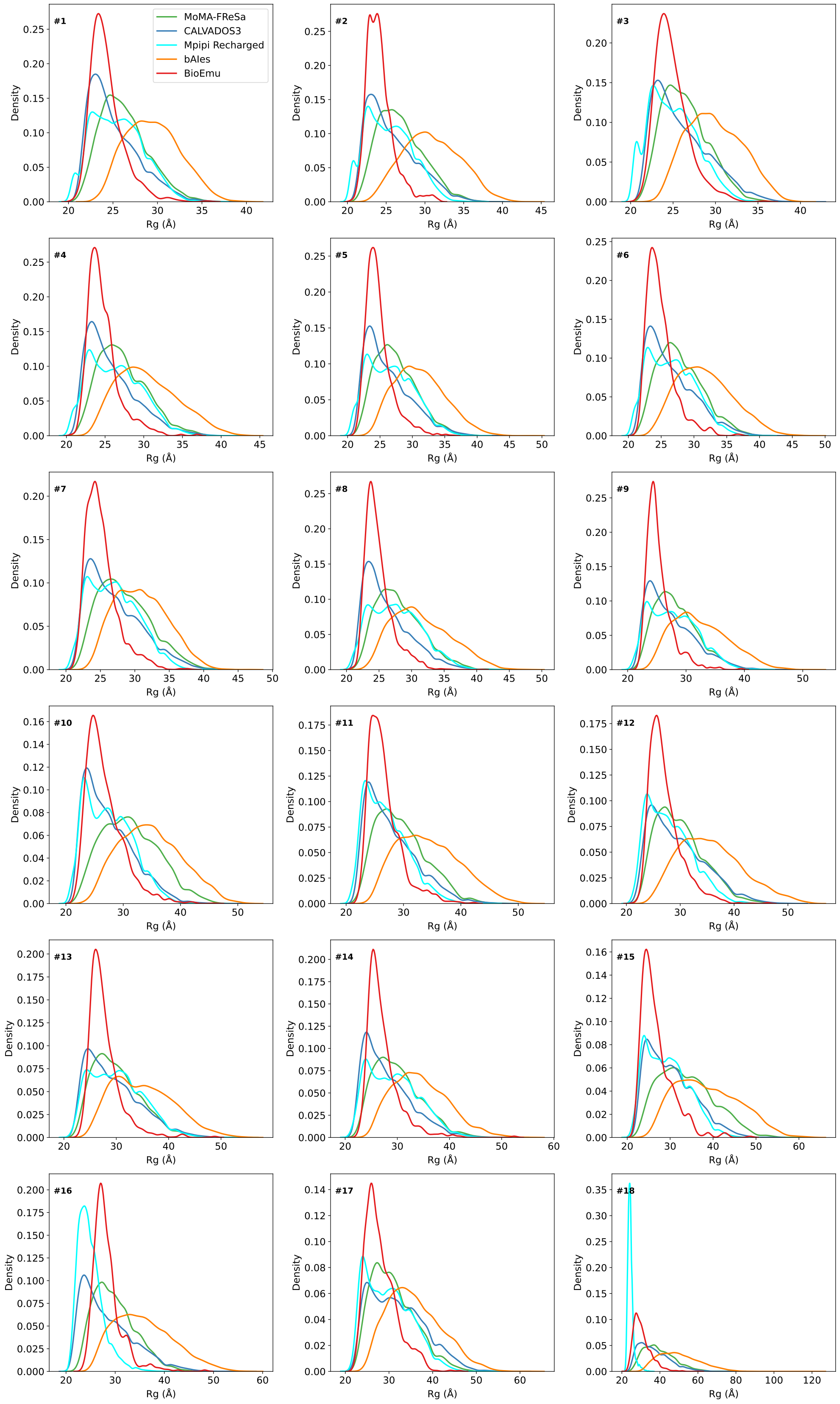


Figure S6. Rg distributions computed from the ensembles generated using the methods tested in this study for the 18 DLD proteins: MoMA-FReSa (green), CALVADOS3 (blue), Mpipi Recharged (cyan), bAIes (orange) and BioEmu (red). Note that the scales of the panels change according to the values obtained. The specific DLD number is shown in each panel.

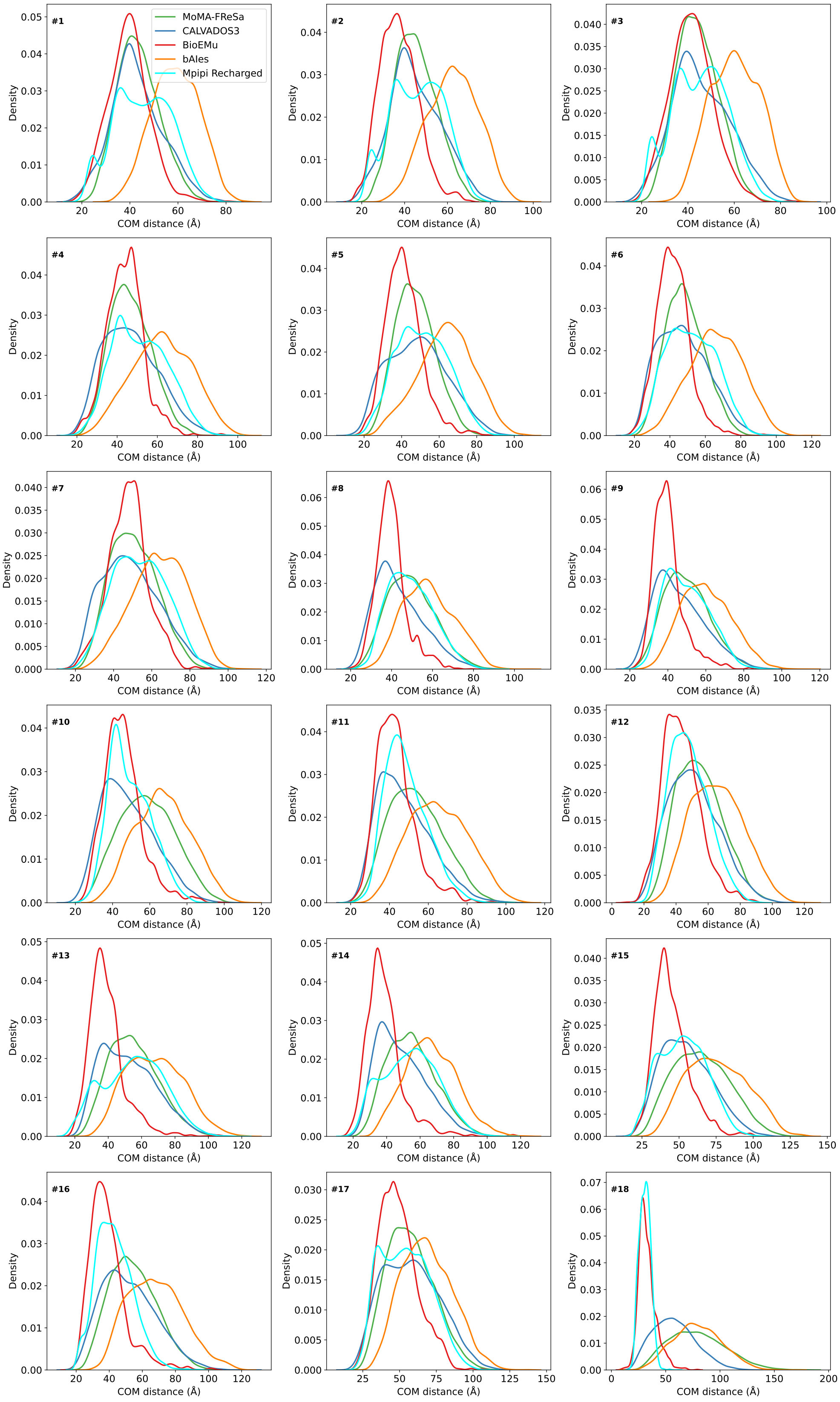


Figure S7. CoM distance distributions computed from the ensembles generated using the methods tested in this study for the 18 DLD proteins: MoMA-FReSa (green), CALVADOS3 (blue), Mpipi Recharged (cyan), bAIes (orange) and BioEmu (red). Note that the scales of the panels change according to the values obtained. The specific DLD number is shown in each panel.

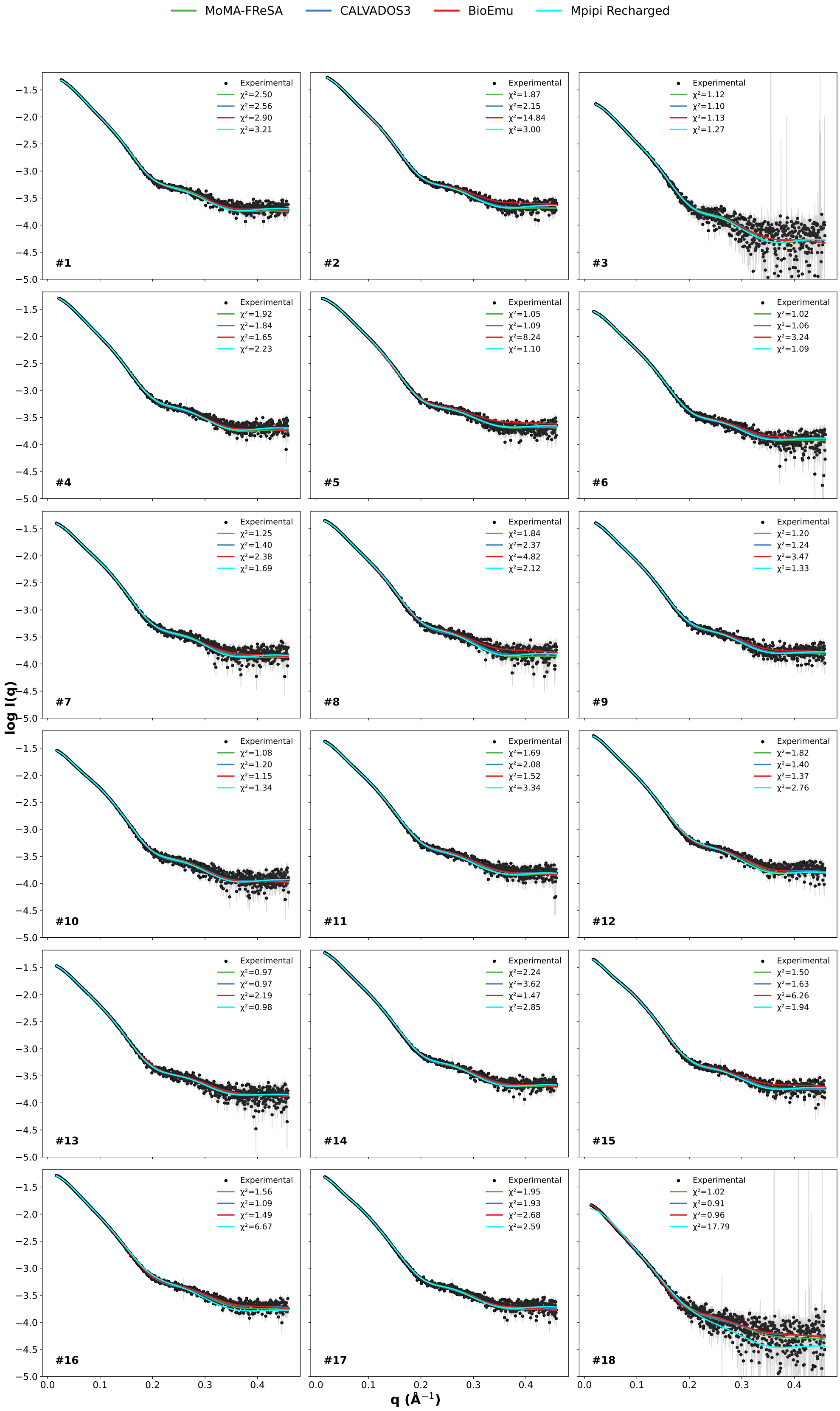


Figure S8. Theoretical scattering profiles fitted against the experimental SAXS curves (in black). MoMA-FReSa is displayed in green, CALVADOS3 in blue, BioEmu in red and Mpipi Recharged in cyan. $\chi^2$-values and the specific DLD number is shown in each panel.

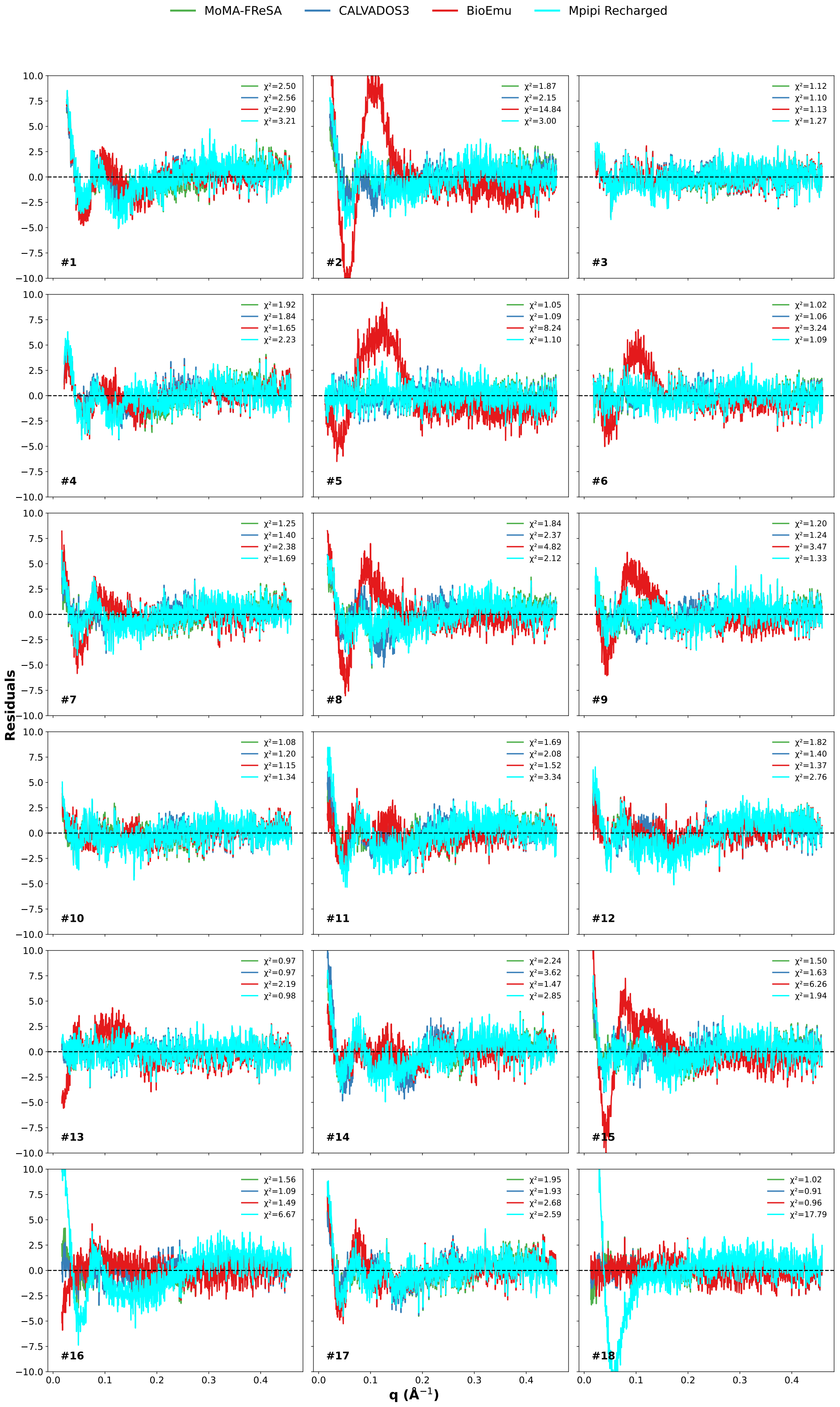


Figure S9. Point-by-point residuals between the SAXS intensity profiles and the fitted theoretical scattering profiles from Figure S8. The specific DLD number is shown in each panel.

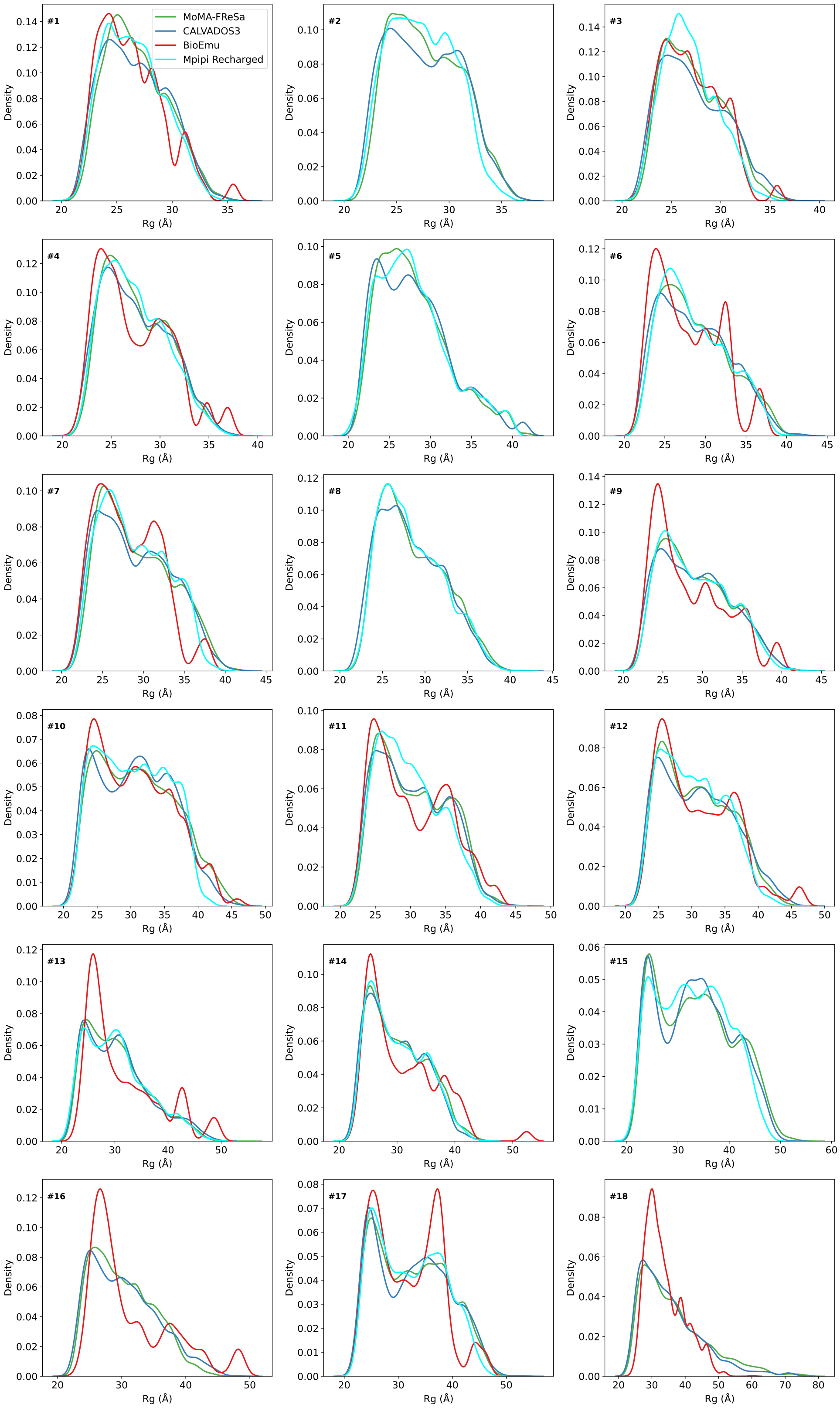


Figure S10. Rg distributions derived from the EOM analyses of the SAXS profile for each DLD protein using the ensembles computed using the different modelling strategies. Note that only distributions obtained with EOM fits with $\chi^2$ values below 4 are shown. This corresponds to MoMA-FReSa (green) and CALVADOS3 (blue) in all cases, BioEMu (red) for 14 cases and and Mpipi Recharged for 16 cases (cyan). The specific DLD number is shown in each panel.

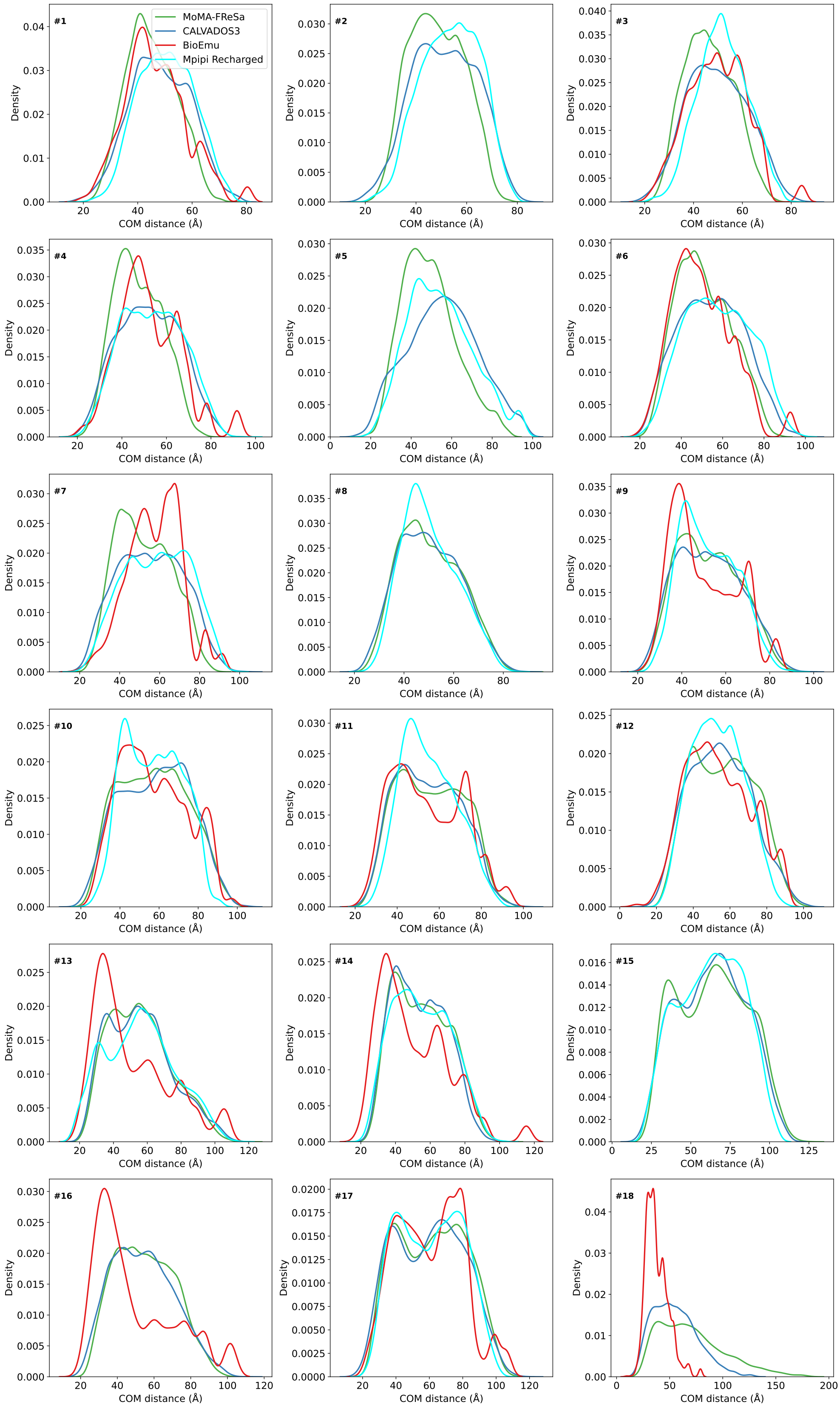


Figure S11. CoM distance distributions computed from the EOM-fitted ensembles for the 18 DLD constructs: MoMA-FReSa (green), CALVADOS3 (blue), BioEmu (red) and Mpipi Recharged (cyan). Note that the scales of the panels change according to the values obtained. The specific DLD number is shown in each panel.

**MGSSHHHHHH**KFTVGNGQNQHKGVNDGFSYEIWLDNTGGNGSMTLGSGATFKAEWNA
AVNRGNFLARRGLDFGSQKKATDYDYIGLDYAATYKQTASASGNSRLCVYGWFQNRGLN
GVPLVEYYIIEDWVDWVPDAQGKMVTIDGAQYKIFQMDHTGPTINGGSETFKQYFSVRQQ
KRTSGHITVSDHFKEWAKQGWGIGNLYEVALNAEGWQSSGVADVTLLDVY**TT**HM**-linker-**
**LESTGC**SVTATRAEEWSDRFNVTYSVSGSSAWTVNLALNGSQTIQASWNANVTGSGSTRT
VTPNGSGNTFGVTVMKNGSSTTPAATCAG**SGGGTA**

His-Tag
GH (PDB: 2C1F)
Cloning site
CBM (PDB:2XBD)

Figure S12. Generic amino acid sequence for the 18 DLD constructs tested in this work. Residues in bold correspond to flexible amino acids, otherwise rigid.
